# Paraxial mesoderm organoids model development of human somites

**DOI:** 10.1101/2021.03.22.436471

**Authors:** Christoph Budjan, Shichen Liu, Adrian Ranga, Senjuti Gayen, Olivier Pourquie, Sahand Hormoz

## Abstract

During the development of the vertebrate embryo, segmented structures called somites are periodically formed from the presomitic mesoderm (PSM), and give rise to the vertebral column. While somite formation has been studied in several animal models, it is less clear how well this process is conserved in humans. Recent progress has made it possible to study aspects of human paraxial mesoderm development such as the human segmentation clock *in vitro* using human pluripotent stem cells (hPSCs), however, somite formation has not been observed in these monolayer cultures. Here, we describe the generation of human paraxial mesoderm (PM) organoids from hPSCs (termed Somitoids), which recapitulate the molecular, morphological and functional features of paraxial mesoderm development, including formation of somite-like structures *in vitro*. Using a quantitative image-based screen, we identify critical parameters such as initial cell number and signaling modulations that reproducibly yielded somite formation in our organoid system. In addition, using single-cell RNA sequencing and 3D imaging, we show that PM organoids both transcriptionally and morphologically resemble their *in vivo* counterparts and can be differentiated into somite derivatives. Our organoid system is reproducible and scalable, allowing for the systematic and quantitative analysis of human spinal cord development and disease *in vitro*.

## Introduction

Paraxial mesoderm (PM) development involves the formation of embryonic segments called somites, which are produced sequentially from the presomitic mesoderm (PSM) and arranged periodically along the anterior-posterior (AP) axis of the vertebrate embryo. Somites give rise and contribute to a variety of tissues including skeletal muscle, dermis, cartilage and bone (Chal and Pourquié, 2017). Somite formation is controlled by a conserved molecular oscillator, the segmentation clock (Dequeant et al., 2006; Hubaud and Pourquié, 2014; Oates et al., 2012). Previous efforts have focused on how this oscillator controls somite formation using a variety of model systems such as mouse, zebrafish, and chick, because of ethical and technical limitations of culturing human embryos. Recently, researchers were able to recapitulate paraxial mesoderm development using human and mouse pluripotent stem cells cultured as 2D monolayers (Chu et al., 2019; Diaz-Cuadros et al., 2020; Matsuda et al., 2020). These cells undergo species-characteristic oscillations similar to their *in vivo* counterparts. However, final stages of somite development and vertebra formation was not observed in currently published human cell culture systems (Palla and Blau, 2020), suggesting that certain aspects of *in vivo* development are not recapitulated in these 2D systems. We reasoned that a 3D cell culture system may exhibit all the stages of PM development including morphogenetic processes associated with somite formation.

Here, we describe an organoid system derived from human iPS cells (hiPSCs) called Somitoids, which faithfully recapitulates functional, morphological and molecular features of paraxial mesoderm development, including formation of somite-like structures *in vitro*. To identify the culture conditions that reproducibly yielded somite formation, we developed a quantitative image-based screening platform for individual organoids. Our screening approach identified the optimal parameter values for the culture conditions such as the initial cell number and the concentration of the chemical modulators. We show using single-cell RNA-sequencing (scRNA-seq), immunofluorescence, and qRT-PCR that Somitoids resemble their *in vivo* counterparts both transcriptionally and morphologically and can be differentiated into somite derivatives such as sclerotome and dermomyotome *in vitro*.

Our Somitoid system is reproducible and scalable, allowing for systematic and quantitative analysis of paraxial mesoderm development and somite formation to study human spinal cord development and disease in a dish. Finally, our approach can be used to systematically screen organoid cultures for desired phenotypes and reproducibility.

## Results

Recently, protocols have been developed to differentiate mouse or human pluripotent stem cells towards paraxial mesoderm using a combination of the WNT agonist CHIR and BMP inhibitor LDN (Chal et al., 2015; Diaz-Cuadros et al., 2020). To adapt the protocol for a 3D model of human somitogenesis, we first optimized the initial conditions of our cultures by generating pluripotent spheroids of defined cell numbers. hiPS cells were allowed to aggregate for 24 hours as suspension cultures in pluripotency media (mTeSR1) in the presence of ROCK inhibitor and Polyvinyl alcohol (PVA) to promote aggregation (Fig 1A). These pluripotent spheroids resemble cavity-stage epiblast embryos as previously described for kidney organoid cultures (Freedman et al., 2015)(Fig 1B and Fig 1-Supplemental Fig 1A). Next, spheroids were cultured in media containing CHIR and LDN (CL), similar as done for the monolayer cultures (Chal et al., 2015; Diaz-Cuadros et al., 2020) but with CHIR at a higher concentration (10 µM). After 24 hours in CL media, epiblast-stage cells transition to a neuromesodermal (NMp) or primitive streak cell fate, characterized by co-expression of T/BRA and SOX2 (Tzouanacou et al., 2009)(Fig 1B). By 48 hours, cells rapidly downregulate SOX2 and express PSM markers, including TBX6 and MSGN1 (Fig 1B). This PSM state persists from day 2 to day 4 and is also characterized by the expression of segmentation clock genes such as HES7 (Fig 1C). On day 5, organoids showed expression of marker genes associated with somite fate as characterized by qPCR (Fig 1C, Fig 1-Supplemental Fig 1B). Taken together, the order of activation of marker genes in the paraxial mesoderm organoids followed the expected stages of differentiation observed during PM development.

**Figure 1.**
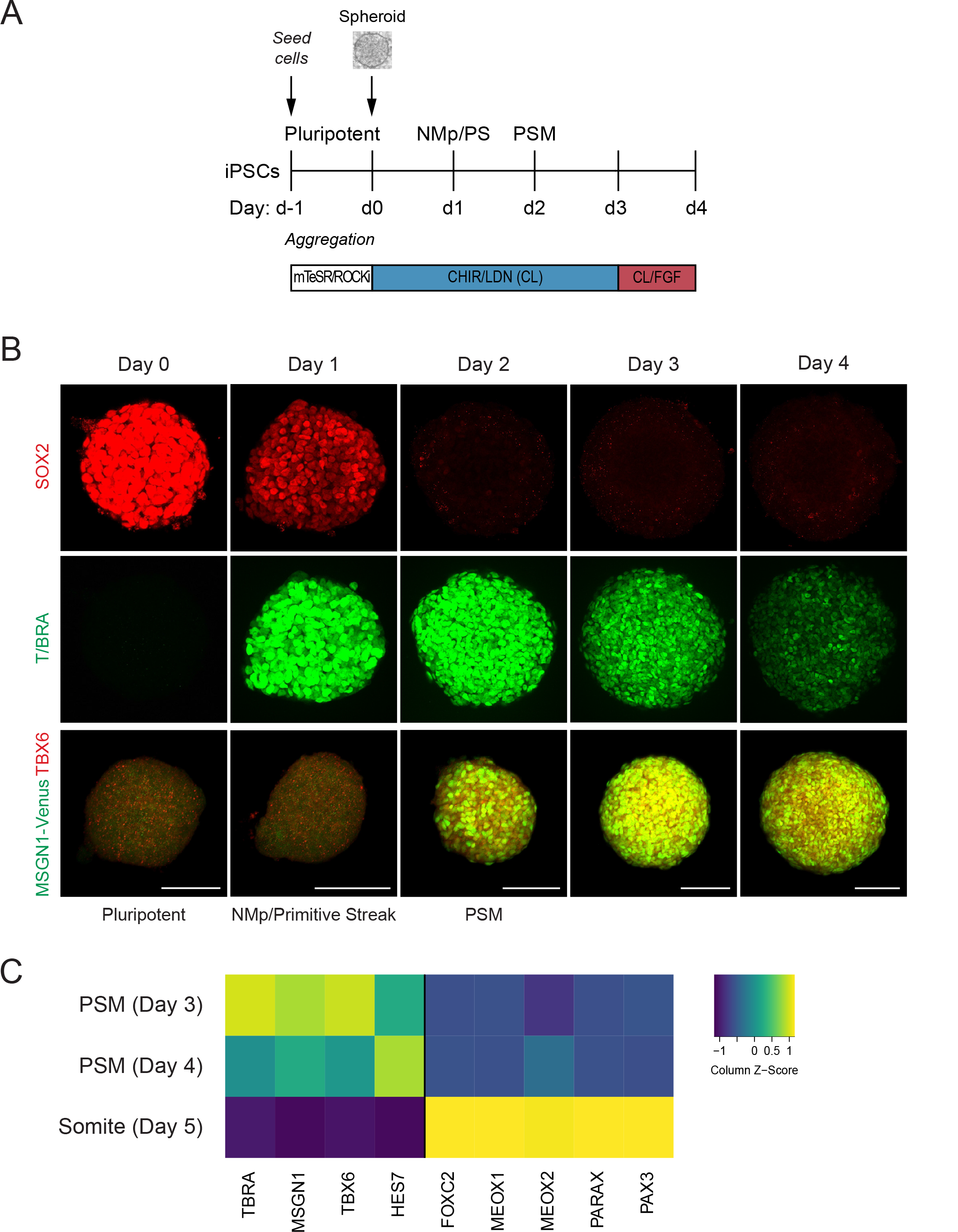
Human PSC-derived paraxial mesoderm organoids turn on marker genes associated with paraxial mesoderm differentiation. (A) Schematic overview of paraxial mesoderm (PM) organoid differentiation protocol from human pluripotent stem cells (PSCs). Human PSCs aggregated and formed spheroids for 24 hours prior to differentiation. For differentiation, spheroids were exposed to Wnt agonist (CHIR) and BMP inhibitor (LDN) for 72 hours. On day 3, FGF2 was added to the media in addition to CHIR and LDN. (B) Immunofluorescence analysis of cell fate-specific marker genes shows progressive differentiation towards PSM fate (top and middle row). Organoids derived from human iPS cells harboring an MSGN1-Venus reporter express TBX6 at the same time as the reporter is activated (bottom row). Scale bar represents 100 µm. Representative images shown from n = 3 independent experiments. Cell lines used: NCRM1 hiPSCs and MSGN1-Venus hiPS reporter cells. (C) qRT-PCR analysis of PSM and Somite markers reveals PSM-to-somite transition from day 4 to day 5. Relative gene expression levels are shown as Z-scores, expressed as fold-change relative to undifferentiated iPS cells (see Methods). Source data is available in Figure 1-Source Data 1.

We observed that following the above protocol resulted in a heterogeneous activation of somite marker genes across cells within the same organoid and across different replicates, as well as a low number of somite-like structures (Figure 1-Supplemental Fig 2). To improve reproducibility, we set out to screen for the optimal initial cell number, the concentration of the signaling factors and the timing of their additions during culture. For our primary screen, we compared organoids with an initial cell number of 500 and 1000 as well as combinations of modulators of candidate signaling pathways that have previously been involved in paraxial mesoderm development and somite formation, including FGF, WNT, BMP and TGF-β (Aulehla et al., 2008; Chal et al., 2015; Hubaud and Pourquié, 2014; Tonegawa et al., 1997; Xi et al., 2017). We primarily focused on the FGF and WNT signaling pathways since their critical role during somitogenesis is well established both *in vivo* (Aulehla et al., 2008, 2003; Delfini et al., 2005; Dubrulle et al., 2001; Dunty et al., 2008; Greco et al., 1996; Yamaguchi et al., 1999) and *in vitro* (Chal et al., 2015; Sakurai et al., 2012; Tan et al., 2013). Furthermore, dual inhibition of FGF and WNT signaling has been used with some success to generate paraxial mesoderm derivatives *in vitro* (Loh et al., 2016; Matsuda et al., 2020). Finally, the BMP and TGF-β signaling pathways have been shown *in vitro* and *in vivo* to have a role in human somitogenesis (Loh et al., 2016; Xi et al., 2017).

**Figure 2.**
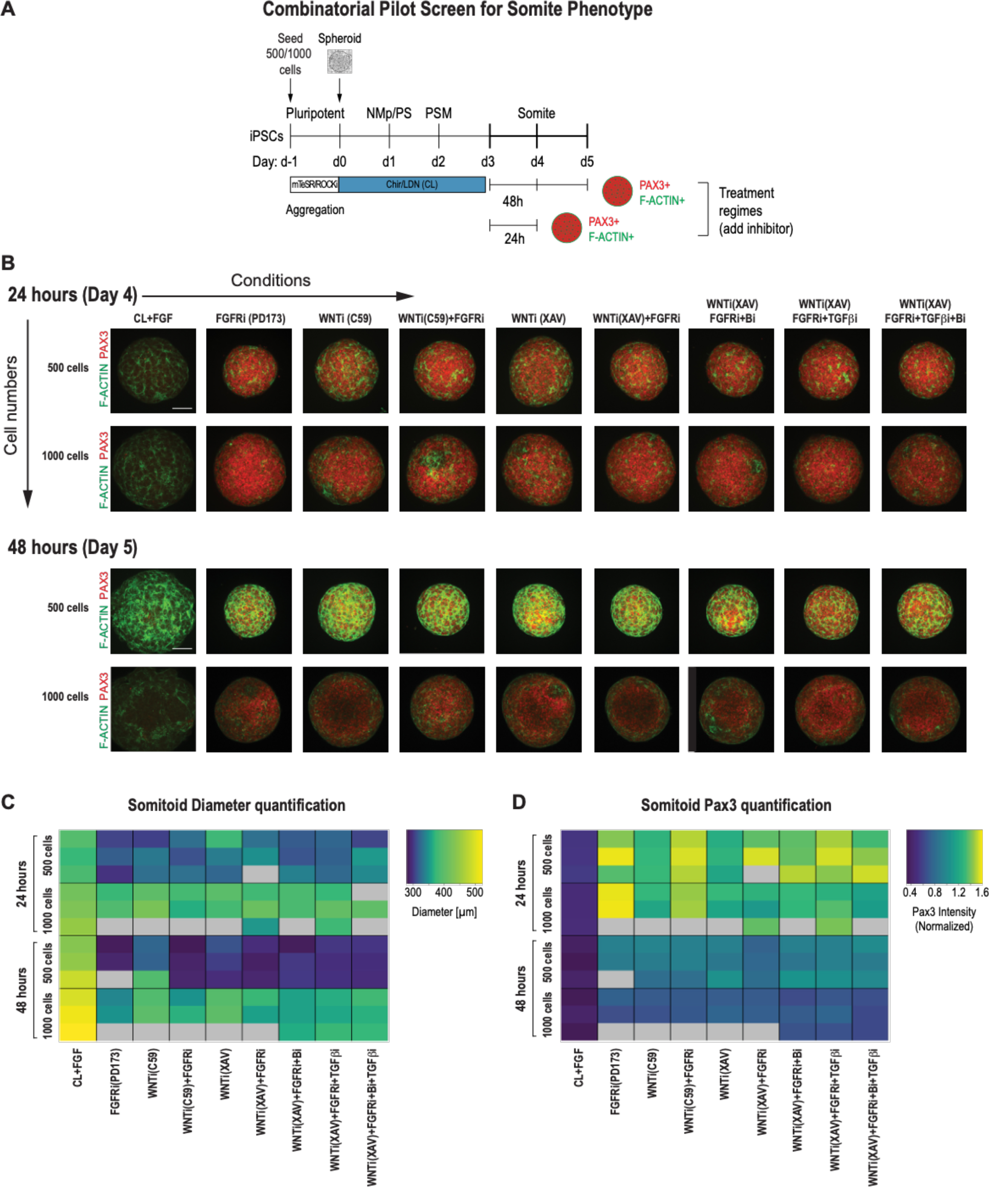
Pilot screen to optimize differentiation conditions for somite phenotype in PM organoids reveals optimal initial number of cells and duration of treatment. (A) Schematic overview of systematic screen in PM organoids (Somitoids). PSM-stage organoids were treated on day 3 for 24 hours or 48 hours with signaling agonists/antagonists. Treated organoids were cultured in basal media with inhibitors as indicated. Control organoids were maintained in CL media with FGF added. NCRM1-derived organoids were used for the screen. (B) Representative immunofluorescent images of day 4 and day 5 organoids after treatment for 24 hours or 48 hours, respectively, stained for somite marker PAX3 and F-ACTIN to visualize rosette-like somite structures. Organoids generated from 1000 cells generally show a more diffuse F-ACTIN pattern compared to organoids made from 500 cells, which exhibit bright foci, consistent with somite formation. Confocal images are shown as maximum intensity z-projections. Scale bar represents 100 µm. Small molecule inhibitors used are indicated in brackets. FGFRi: FGF receptor inhibitor (PD173074). WNTi: Wnt inhibitor (C59 or XAV939). Bi: BMP inhibitor (LDN). TGF-βi: TGF-β inhibitor (A-83-01). Representative image shown for each condition from 3 organoid replicates. (C) Automated quantification of organoid diameter for each organoid/replicate treated as indicated (see Methods for details). Three organoids per condition were characterized except where indicated with grey boxes. Organoids initiated from 500 cells show a decreased diameter when treated for 48 hours compared with 24 hours. Source data is available in Fig 2-Source Data 1. (D) Automated quantification of normalized average PAX3 intensity for each organoid/replicate treated as indicated. Three organoids are shown per condition except where indicated with grey boxes. Organoids initiated from both 500 and 1000 cells show higher average normalized PAX3 levels when treated for 24 hours compared with 48 hours. Source data is available in Fig 2-Source Data 2.

PSM-stage organoids on day 3 were treated with signaling modulators for 24 and 48 hours, and somite fate and morphogenesis was assessed using PAX3, a somite fate marker, and F-ACTIN, a structural marker of somite formation (Fig 2A). We chose day 3 as a starting point for our systematic screen because PSM marker gene expression was more uniform compared to day 2 organoids (Fig 1B), and day 4 organoids are not significantly different from day 3 organoids based on a previous study and our own immunostaining and qPCR data (Figure 1B,C and Diaz-Cuadros et al., 2020). To quantitatively compare conditions, we developed an image analysis pipeline to determine organoid diameter and normalized average PAX3 expression intensity per organoid in an automated manner (Fig 2C,D and Fig 2-Supplemental Fig 2B). We analyzed three organoids per condition. Strikingly, all organoids that were treated with any combination of FGF or WNT inhibitor reproducibly expressed PAX3 within 24 hours of treatment (Fig 2B,D). However, organoids that were initiated from 1000 cells displayed a higher fraction of PAX3-negative cells compared with organoids initiated from 500 cells, even though the average PAX3 expression levels across the entire organoid was comparable (Figure 2D and Figure 2-Supplemental Fig 2A). Additional staining for SOX2, a neural marker, showed that PAX3-negative cells expressed SOX2, suggesting that our paraxial mesoderm organoids derive from neuro-mesodermal progenitors (Fig 1B and Fig 2-Supplemental Fig 2A). In addition, staining for F-ACTIN, together with nuclear expression of PAX3, more consistently revealed somite-like structures (radial arrangement of PAX3+ columnar cells with expression of apical F-ACTIN in the central cavity) in organoids made from 500 cells compared with organoids made from 1000 cells (Figure 2B and Figure 2-Supplemental Fig 3A,B). Taken together, based on these observations, we used 500 cells as the initial cell number going forward.

**Figure 3.**
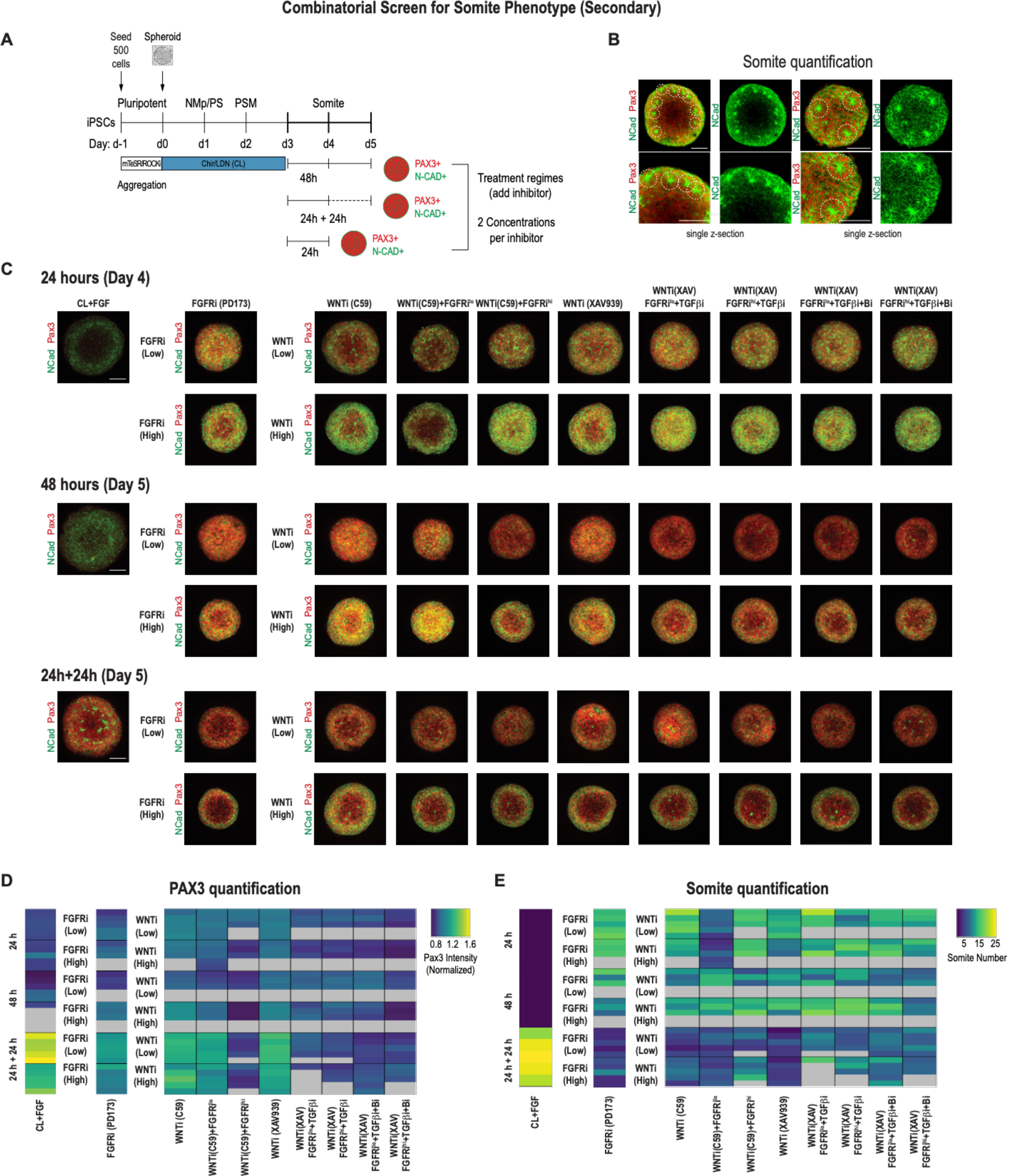
Secondary screen of PM organoids identifies optimal differentiation protocol for somite formation. (A) Schematic overview of secondary screen in PM organoids. PSM-stage organoids were treated on day 3 for 24 hours followed by measurement, 24 hours of treatment followed by 24 hour culture in basal media (no added factors) and then measured (24h+24h), and 48 hours of treatment followed by measurement. Treated organoids were cultured in basal media with inhibitors as indicated. WNT and FGF inhibitors were tested at two different concentrations. Control organoids were maintained in CL media with FGF added. NCRM1-derived organoids were used for the screen. (B) Representative immunofluorescent images of day 5 organoids stained for somite markers PAX3 and NCAD showing the rosette-like structures which were scored as somites based on expression of somite fate markers and structural features (scoring criteria detailed in Methods section). Images are shown as individual z-sections. (C) Representative immunofluorescent images of day 4 and day 5 organoids stained for somite markers PAX3 and NCAD to visualize rosette-like somite structures. Confocal images are shown as maximum intensity z-projections. Scale bar represents 100 µm. Small molecule inhibitors used are indicated in brackets. FGFRi: FGF receptor inhibitor (PD173074). WNTi: Wnt inhibitor (C59 or XAV939). Bi: BMP inhibitor (LDN). TGF-βi: TGF-β inhibitor (A-83-01). Representative image shown for each condition from 5 organoid replicates. (D) Automated quantification of normalized average PAX3 intensity for each organoid/replicate treated as indicated (see Methods for details). Five organoids are shown per condition except as indicated with grey boxes. Several inhibitor combinations with a treatment regime of 24 hour treatment followed by 24 hour cultured in basal media show highest average PAX3 levels. Source data is available in Fig 3-Source Data 1. (E) Quantification of the number of somite-like structures for each condition. Each row represents one organoid replicate. Five organoids are shown per condition except where indicated with grey boxes. Organoids which were maintained in CL media with added FGF for 24 hours followed by culture in basal media for 24 hours reproducibly exhibit the highest number of somites per organoid. Source data is available in Fig 3-Source Data 2.

Organoids that were treated for 48 hours with signaling pathway modulators showed an overall decrease of PAX3 expression compared with organoids treated for only 24 hours across replicates, indicating that prolonged signaling manipulation does not improve the somite phenotype (Fig 2D). Additionally, organoids initiated from 500 cells and then treated for 48 hrs had a smaller diameter compared with organoids of the same initial cell number that were treated for only 24 hours (Fig 2C). This suggests that long-term inhibition of WNT and/or FGF, known mitogenic signaling pathways, has detrimental effects on proliferation or cell survival. These results indicate that treatment of PSM-stage organoids with pathway modulators for 24 hours is sufficient to induce somite fate.

Next, we set out to optimize the culture conditions to increase the number of somite-like structures in addition to the expression levels of the somite marker genes. We looked for morphological hallmarks of somite formation, specifically the formation of rosette-like structures consisting of radially arranged bottle-shaped PAX3+ epithelial cells with their NCAD+ apical surface facing a central cavity (Figure 3-Supplemental Fig 1A,B and Figure 3-Video 1).

We used 500-cell spheroids as an initial starting point and compared two different inhibitor doses for FGF and WNT in addition to the other pathway modulators applied to PSM-stage organoids on day 3. We characterized the organoids after treating them for 24 hours, 48 hours, and 24 hours followed by culture in basal media without any added factor for an additional 24 hours (5 organoids per condition, Fig 3A). In addition to quantifying PAX3 levels (Fig 3D, Fig 3-Supplemental Fig 2B), we also counted the number of somite-like structures per organoid (Fig 3B, Fig 3E, Fig 3-Supplemental Fig 2A,B; see Methods section for a description of the scoring criteria). Comparing PAX3 expression levels in our treated organoids, we observed that somite fate can be broadly induced across a range of treatment regimes, concentrations and types of WNT and/or FGF inhibitors (Fig 3C,D and Fig 3-Supplemental Fig 2C). However, the number of somite-like structures is not necessarily correlated with average PAX3 expression levels. For example, the numbers of somite-like structures in several conditions (FGFRi^hi^/PD173, WNTi^hi^/C59, WNT^hi^/C59+FGFRi^lo^, WNTi^hi^/XAV) were lower in organoids treated for 24 hours followed by culture in basal media for 24 hours compared with organoids that were treated with the same inhibitors for 48 hours, even though they exhibited higher PAX3 expression levels on average (Fig 3D,E and Fig 3-Supplemental Fig 2A). This suggests that marker gene expression alone may not be the best predictor when screening for morphologically complex phenotypes such as somite formation.

Surprisingly, organoids which were cultured for an additional 24 hours (day 3 to day 4) in FGF, WNT pathway agonist and BMP inhibitor, considered a treatment control, followed by culture in basal media only for another 24 hours, consistently exhibited the highest number of somite-like structures across all organoid replicates (Fig 3E, Figure 3-Supplemental Fig 1A-B, Fig 3-Supplemental Fig 2A-B and Figure 3-Video 1) as well as technical replicates (Figure 3-Supplemental Fig 3A). Additionally, the average PAX3 expression was among the highest of all conditions tested (Fig 3D). This suggests that simply removing FGF/WNT pathway agonists and BMP inhibitor, which maintain cells in a PSM state, is sufficient to reproducibly induce somite fate and morphological formation of somite-like structures (Fig 3-Supplemental Fig 2A, Fig 3-Supplemental Fig 3A). Computing the variation of somite numbers across the 5 organoids confirmed that this phenotype was highly reproducible (coefficient of variation = 11.1%; Fig 3-Supplemental Fig 2B). Finally, our optimized protocol reproducibly yielded efficient somite induction in multiple genetically independent hiPS cell lines (Figure 3-Supplemental Fig 3B). Taken together, we determined that initiating the protocol with 500 cells and treating day 3 organoids with CL+FGF for 24 hours followed by culture in basal media for an additional 24 hours yields the most robust somite induction while minimizing variation between experiments (technical variation) and different cell lines (biological variation). This optimized differentiation protocol was therefore used for all subsequent experiments.

To further characterize the developmental trajectory and transcriptional states of our Somitoid system, we collected 15,558 cells (after post-processing) over the course of the optimized five day differentiation protocol at time-points that capture the key transition steps (day 1, day 2, day 3, day 5) and performed single-cell RNA sequencing (scRNA-seq; Fig 4A). Multiple organoids were used to obtain the required number of cells at each timepoint (see Methods). We first combined all the cells across the time-points and clustered them using the Leiden clustering algorithm (Traag et al., 2019). Predominantly, the 4 major clusters corresponded to cells from the 4 different time-points. Therefore, cells at each time-point have transcriptional states that are different compared with cells from the other time-points. In addition, within each time-point, cells exhibit similar transcriptional profiles as indicated by the uniformity of the expression levels of marker genes across individual cells (Fig 4E, Fig 4-Supplemental Fig 1,2,3).

**Figure 4.**
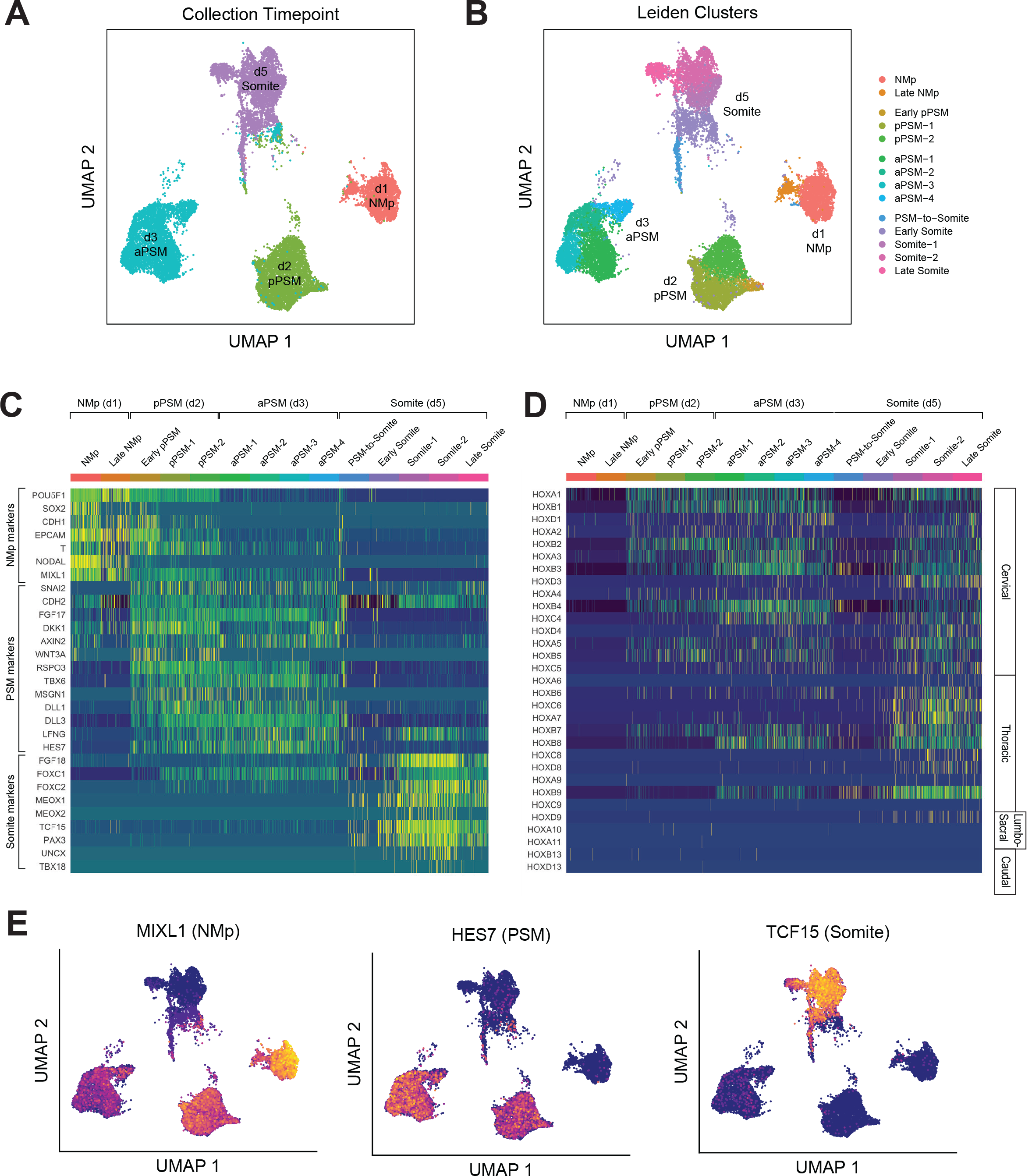
Single-cell RNA sequencing analysis of PM organoids (Somitoids) reveals differentiation trajectory from NMp-like cells to somite-stage paraxial mesoderm. (A) UMAP (Uniform Manifold Approximation and Projection) of single-cell transcriptomes of differentiating human PM organoids, colored by collection time point (15,558 cells). NCRM1-derived organoids were used to collect single cells. (B) UMAP of human PM organoids, colored by assigned Leiden cluster identity based on marker gene expression profile (see Methods). (C) Heat map of selected marker genes of paraxial mesoderm differentiation. Collection time point and Leiden cluster identities are indicated. Marker genes are grouped based on primary associated cell fate as indicated. (D) Heat map of single-cell HOX gene expression levels. Cells are grouped by Leiden cluster identity. Hox genes are ordered by position, with anatomical positions of HOX paralogues indicated on the right. (E) UMAP plots overlaid with normalized transcript counts of representative cell fate marker genes.

Cells collected on day 1 exhibited gene expression profiles similar to primitive streak or neuro-mesodermal progenitors, expressing genes such as SOX2, T/BRA, MIXL1 and NODAL (Fig 4C,E, Fig 4-Supplemental Fig 1). Starting on day 2, cells expressed canonical PSM marker genes such as TBX6, MSGN1, WNT3A, RSPO3 and clock genes of the Notch signaling pathway including HES7, LFNG, DLL1 and DLL3 (Fig 4C,E; Fig 4-Supplemental Fig 2). Day 5 cells expressed somite-marker genes such as TCF15, PAX3, FOXC2 and MEOX2 (Fig 4C,E; Fig 4-Supplemental Fig 3). Interestingly, a subset of the day 5 cells also expressed somite polarity markers, UNCX and TBX18, which suggests faithful recapitulation of somite patterning in Somitoids (Fig 4C; Fig 4-Supplemental Fig 3). Furthermore, two of the sub-clusters (‘PSM-to-Somite’, ‘Early Somite’), which are characterized by co-expression of PSM and somite marker genes, were comprised of both day 3 and day 5 cells, indicating that the PSM-to-somite transition is captured in our *in vitro* system (Fig 4A,B). Interestingly, one somite sub-cluster (‘Late Somite’) was enriched for myogenic genes (MYL4/6/7/9, TROPONIN L1) and sclerotome genes (TWIST1, COL1A1, COL11A1, COL7A1, ACTA2), suggesting that these cells represent more downstream fates of somite-derived cells (Fig 4B; Fig 4-Supplemental Fig 4A). Finally, we observed the expected sequential activation pattern of the HOX genes in our Somitoid system starting with HOXA1 on day 1, followed by other cervical and thoracic HOX genes on day 2-3, to HOXD9, a lumbosacral HOX gene, in the somite-stage organoids (day 5; Fig 4D and Fig 4-Supplemental Fig 4B). Taken together, our scRNA-seq analyses show that our Somitoid system faithfully recapitulates the gene regulatory programs of human paraxial mesoderm development and generates mature somite cells, which express the full repertoire of known marker genes. Furthermore, we did not detect cells of different developmental origins, suggesting that we are generating homogeneous organoids containing only paraxial mesoderm derivatives.

We next assessed whether our *in vitro* derived somites show similar spatial organization and size distribution to their *in vivo* counterparts. To independently confirm some of the somite marker genes that we identified in our scRNA-seq data set, we measured expression levels of TCF15/PARAXIS, PAX3, and F-ACTIN in day 5 Somitoids using whole-mount immunostaining (Fig 5A). Day 5 cells co-expressed both somite marker genes TCF15 and PAX3 throughout the Somitoid and somite-like structures displayed apical localization of F-ACTIN. To determine whether *in vitro*-derived somites were similar in size to human embryonic segments, we compared them with Carnegie stage 9-11 early human somites (Fig 5B; see Methods section for a description of how somite sizes were quantified). Organoid-derived somites were similar in size (median area = 8892 µm^2^, interquartile range/IQR = 7698-10682 µm^2^) to Carnegie stage 11 somites (median area = 9681 µm^2^, IQR = 8262-11493 µm^2^) but larger than earlier-stage human somites (Carnegie stage 9 somites, median area = 4399 µm^2^, IQR = 4089-4433 µm^2^; Carnegie stage 10 somites, median area = 4704 µm^2^, IQR = 4477-5343 µm^2^; see Fig 5B). Together, these results suggest that our organoid-derived somites share spatial and molecular features as well as overall size with their *in vivo* counterparts.

**Figure 5.**
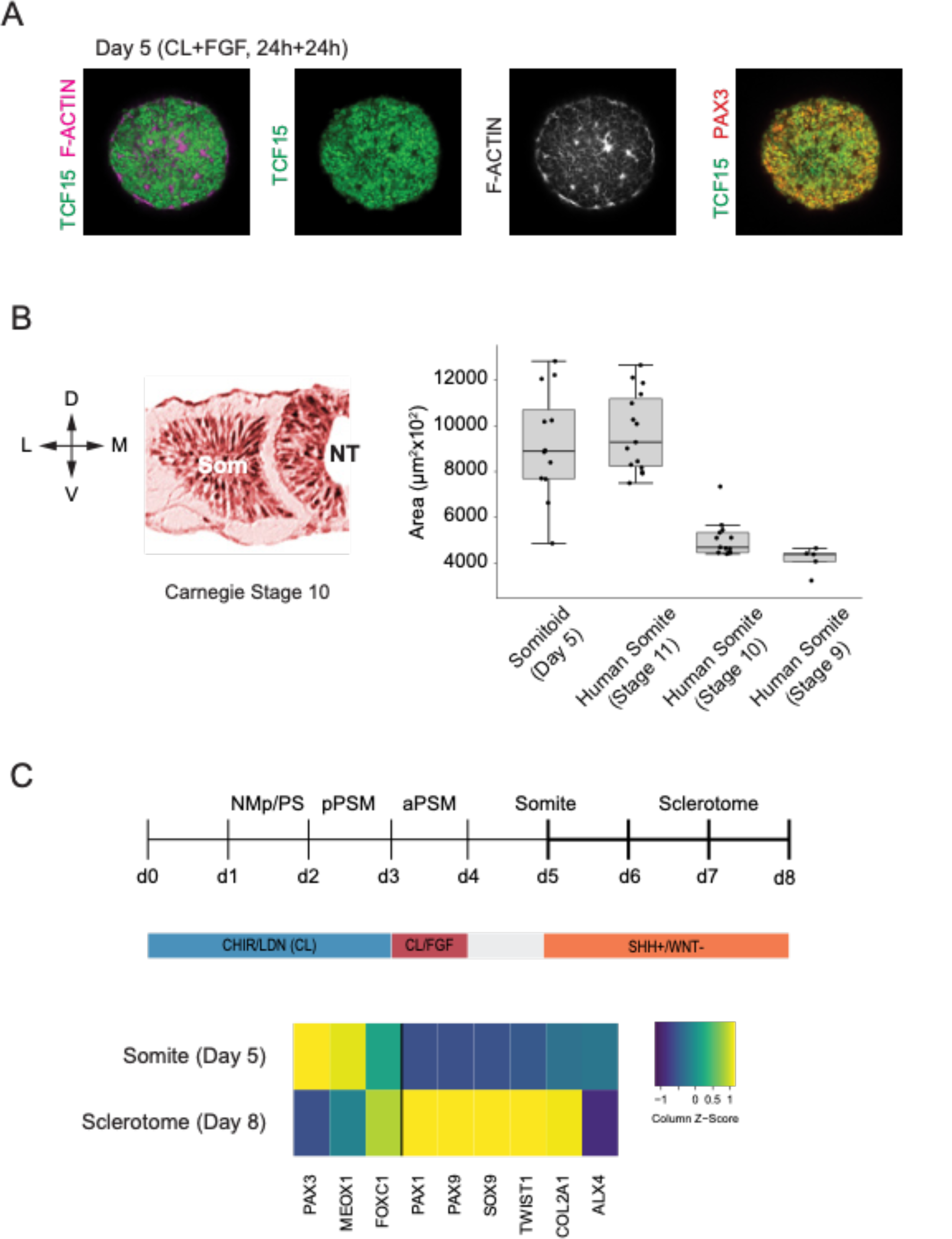
Somitoids express known somite-stage specific marker genes and can differentiate to sclerotome fate. (A) Whole-mount Immunofluorescence analysis of day 5 Somitoids reveals co-expression of somite markers TCF15/PARAXIS and PAX3 and polarized rosette-like structures as indicated by F-ACTIN localization, suggesting that somite-like structures resemble *in vivo* counterparts in both a molecular and morphological manner. Representative images are shown as maximum intensity z-projections from 3 organoid replicates. NCRM1 hiPS cells were used to generate Somitoids. (B) Quantification of somite sizes in day 5 human Somitoids and human embryos (Carnegie stage 9, 10 and 11) reveals that the median and interquartile range of *in vitro* somites sizes (calculated as area) is comparable to Carnegie stage 11 human somites *in vivo* (see Methods section). Boxes indicate interquartile range (25th percentile to 75th percentile). End of whiskers indicate minimum and maximum. Points indicate individual somites. Central lines represent the median. Carnegie embryo data were obtained from the Virtual Human Embryo Project (https://www.ehd.org/virtual-human-embryo). NCRM1 hiPS cells were used to generate Somitoids. Source data is available in Fig 5-Source Data 1. (C) Sclerotome differentiation of Somitoids. Day 5 Somitoids were exposed to SHH agonist and WNT inhibitors to induce sclerotome differentiation as indicated. qPCR analysis of somite and sclerotome markers reveals induction of sclerotome markers on Day 8. Relative gene expression levels are shown as Z-scores, expressed as fold-change relative to undifferentiated iPS cells (see Methods). NCRM1 hiPS cells were used to generate Somitoids. Source data is available in Fig 5-Source Data 2.

Finally, we assessed whether Somitoids can give rise to downstream paraxial mesoderm derivatives of sclerotome and dermomyotome. First we differentiated Somitoids to sclerotome by exposing day 5 organoids to SHH agonist and WNT inhibitor to mimic the signaling environment of ventral somites *in vivo* (Fan et al., 1995; Fan and Tessier-Lavigne, 1994; Loh et al., 2016). After 3 days, the organoids showed a robust induction of canonical sclerotome marker genes such as PAX1, SOX9 and COL2A1 (Fig 5C). Additionally, day 5 Somitoids were differentiated towards dermomyotome by exposing them to WNT/BMP agonists and SHH inhibitor for 48 hours followed by dissociation and culture on Matrigel as a monolayer in muscle differentiation medium (Loh et al., 2016; Matsuda et al., 2020) to further differentiate them to skeletal muscle. Immunostaining for Myosin heavy chain (MYH1, a myocyte/myotube marker) confirmed that our Somitoid-derived cells can generate skeletal muscle derivatives *in vitro* (Figure 5-Supplemental Fig 1). These data demonstrate that our *in vitro* induced somites maintain their ability to differentiate further into somitic mesoderm derivatives of the sclerotome and dermomytome lineages.

## Discussion

Here, we reported the generation of human paraxial mesoderm organoids from hPSCs that reproduce important features of somitogenesis not previously captured in conventional monolayer differentiation cultures, most notably somite formation. Using a simple suspension culture which does not require manual matrix embedding, we identified optimal differentiation conditions by systematically screening initial cell numbers and modulating the signaling factors. Importantly, our culture conditions are compatible with high-throughput screening approaches. Many established organoid protocols currently have limited applications because they are not reproducible. Therefore, we set out to identify the optimal differentiation conditions that minimized the variability between organoids as quantified by automated image analysis.

One critical parameter we identified in our screens was the initial cell number used for aggregation. Our results suggest that if the initial cell number is above a certain threshold then somite fate cannot be induced in a homogeneous manner in our organoid system. This result is in line with previous findings in 3D models such as gastruloids, multi-axial self-organizing aggregates of mouse ES cells, which exhibit a higher degree of variability and multiple elongations when the number of initial cells exceeds a threshold (Beccari et al., 2018; van den Brink et al., 2014). Another important finding of our screens was that simply removing FGF, WNT pathway agonist as well as BMP inhibitor yielded the most reproducible and efficient somite-forming organoids. This treatment regime does not necessarily follow from applying prior *in vivo* and *in vitro* knowledge of somitogenesis. Previous protocols have used FGF and WNT inhibitors (Matsuda et al., 2020) or inhibition of all four candidate signaling pathways (FGF, WNT, BMP and TGF-β) to induce somite fate (Loh et al., 2016) in monolayer cultured hiPSCs. While these conditions similarly induced somite fate marker genes in our 3D system, removal of FGF/WNT agonists and BMP inhibitor overall performed better as indicated by larger organoid diameters, higher average PAX3 expression levels, and higher number of somite-like structures. In line with these findings, our single-cell RNA-seq analysis revealed that day 5 somite-like cells from our optimized protocol autonomously downregulate WNT target genes (DKK1, AXIN2, WNT3A, RSPO3; Fig 4-Supplemental Fig 2) and FGF target genes (FGF8, FGF17, SPRY4, DUSP6/MKP3, SEF/IL17RD; Fig 4-Supplemental Fig 5). Finally, our screening results also suggest that focusing on marker gene induction as a phenotypic readout alone is not sufficient to optimize culture conditions of more complex organoid models such as somitogenesis.

The single-cell RNA-seq analysis of our hPSC-derived Somitoids independently confirmed our immunostaining and qRT-PCR results and showed that all major paraxial mesodermal cell types are generated, consistent with the cell types observed during PM development. Comparing our single-cell dataset with previously published *in vitro* generated human PM transcriptomic data of monolayer cultures (Diaz-Cuadros et al., 2020; Matsuda et al., 2020) reveals a similar pattern of activation of marker genes. Diaz-Cuadros and colleagues did not generate bona fide somitic cells as their final time point cell population does not express canonical somitic markers. Matsuda et al. indeed show expression of several somitic marker genes including TCF15, MEOX1, and PAX3 based on bulk RNA-seq data. Interestingly, our own analysis of day 5 somitic cells revealed multiple distinct sub-clusters, suggesting transcriptional heterogeneity within this population, which could have not been inferred from bulk data (Figure 4B,C and Figure 4-Supplemental Fig 4). Importantly, neither of these papers report formation of somite-like structures, suggesting that transcriptional similarity alone is not sufficient to predict morphological features, in line with our screening results showing that average marker gene expression is not a good predictor of *in vitro* somite induction efficiency (Fig 3D,E and Figure 3-Supplemental Fig 2A-C).

While expression patterns of canonical somitic marker genes seem to be conserved in humans, it will be interesting to perform detailed gene expression analysis to identify putative human-specific genes of somite differentiation. Since our Somitoid system is reproducible, it could serve as a versatile platform to perform functional screens of human-specific or disease-relevant genes using CRISPR/Cas9 or small molecule inhibitor libraries. Somitoids thus provide a powerful *in vitro* system for studying the regulation and dynamics of human somitogenesis, including somite formation.

One limitation of this work is that the current protocol does not produce an anterior-posterior axis and therefore does not generate somites in a bilaterally symmetric fashion as in the vertebrate embryo. A similar phenotype was recently reported in mouse gastruloids that were embedded in matrigel to promote self-organization into trunk-like structures (Veenvliet et al., 2020). Chemical modulation of BMP and WNT signaling pathways in matrigel-embedded gastruloids resulted in formation of somites arranged like a bunch of grapes, similar to what we observed in our system. In standard culture conditions, gastruloids recapitulate the axial organization of the embryo, which is missing in our Somitoids (Beccari et al., 2018; Moris et al., 2020). To expand the patterning and morphogenetic potential of our Somitoid system, our approach could be combined with a microfluidics setup to generate spatio-temporally controlled morphogen gradients (Manfrin et al., 2019). In summary, Somitoids provide a scalable, reproducible and easy to manipulate platform to study molecular networks underlying the differentiation of paraxial mesoderm, as well as the morphogenetic processes of somite formation. Furthermore, Somitoids represent a promising *in vitro* system to study congenital diseases that are linked to the human segmentation clock and somite formation, such as congenital scoliosis.

## Supporting information

Supplemental Table 1, cluster-based marker genes of scRNA-seq dataset

Supplemental Table 2, RT-qPCR primer sequences

Figure 3-Video 1, confocal z-stacks of Somitoids and control organoids immunostained for PAX3 and NCAD showing in vitro somites

## Acknowledgements

This work was supported by the National Institutes of Health (NIH) P01 Pilot Grant (P01GM099117) to CB and SH. CB and SH also acknowledge funding from NIH NHLBI (R01HL158269) and Chan Zuckerberg Initiative (CZI) funding for Collaborative Computational Tools for a Human Cell Atlas (2018-183143). We would like to acknowledge support of the Nikon Imaging Center at Harvard Medical School for image acquisition and consulting.

## Author Contributions

C.B. designed, optimized and performed the experiments. C.B. analyzed the experimental and single-cell RNA-seq data with help from S.G. S.L. assisted with single-cell sample collection and sequencing library preparation. A.R. contributed to an early version of the differentiation protocol. C.B. and S.H. wrote the paper. O.P. and S.H. supervised the project.

## Materials and Methods

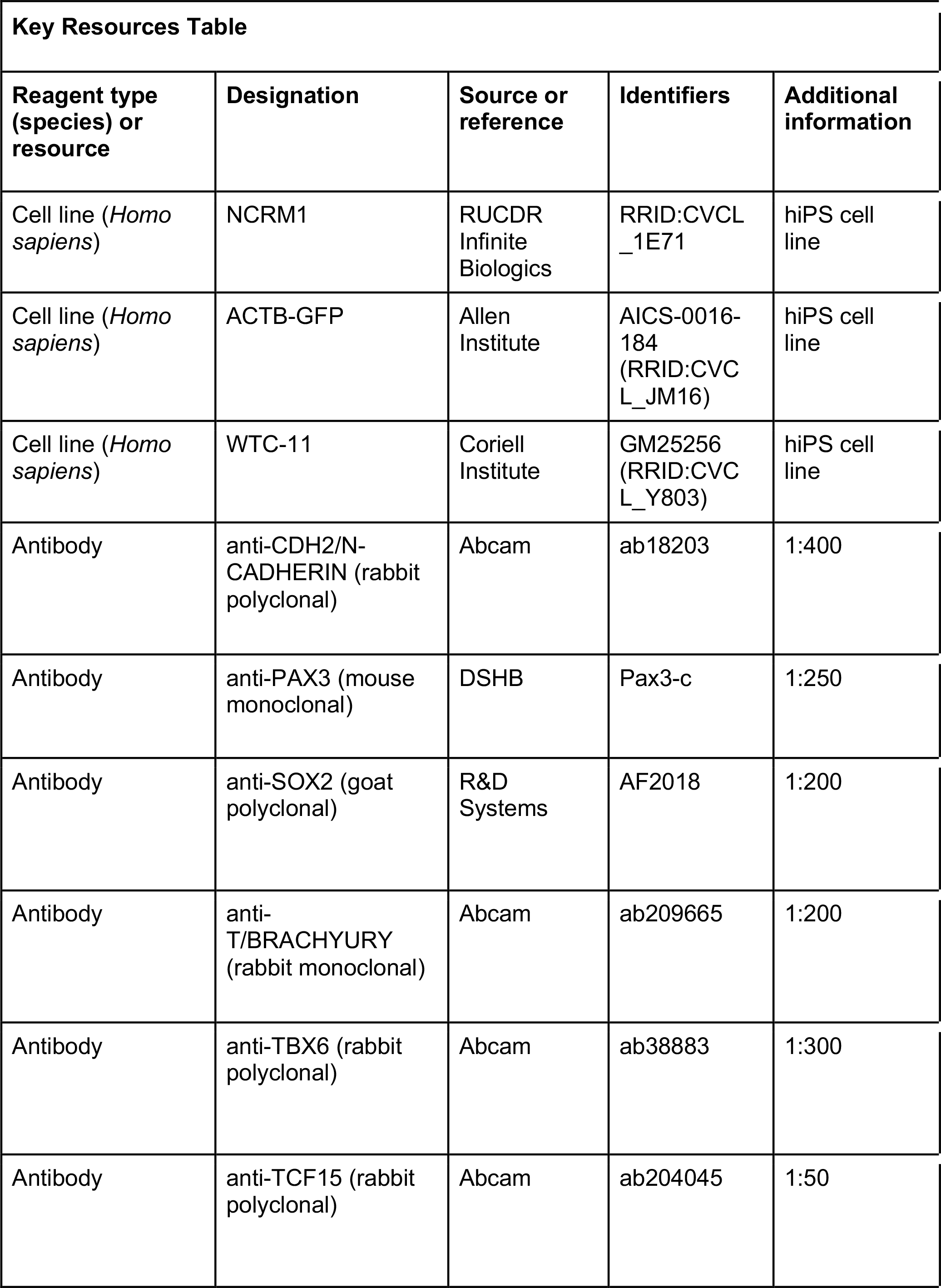
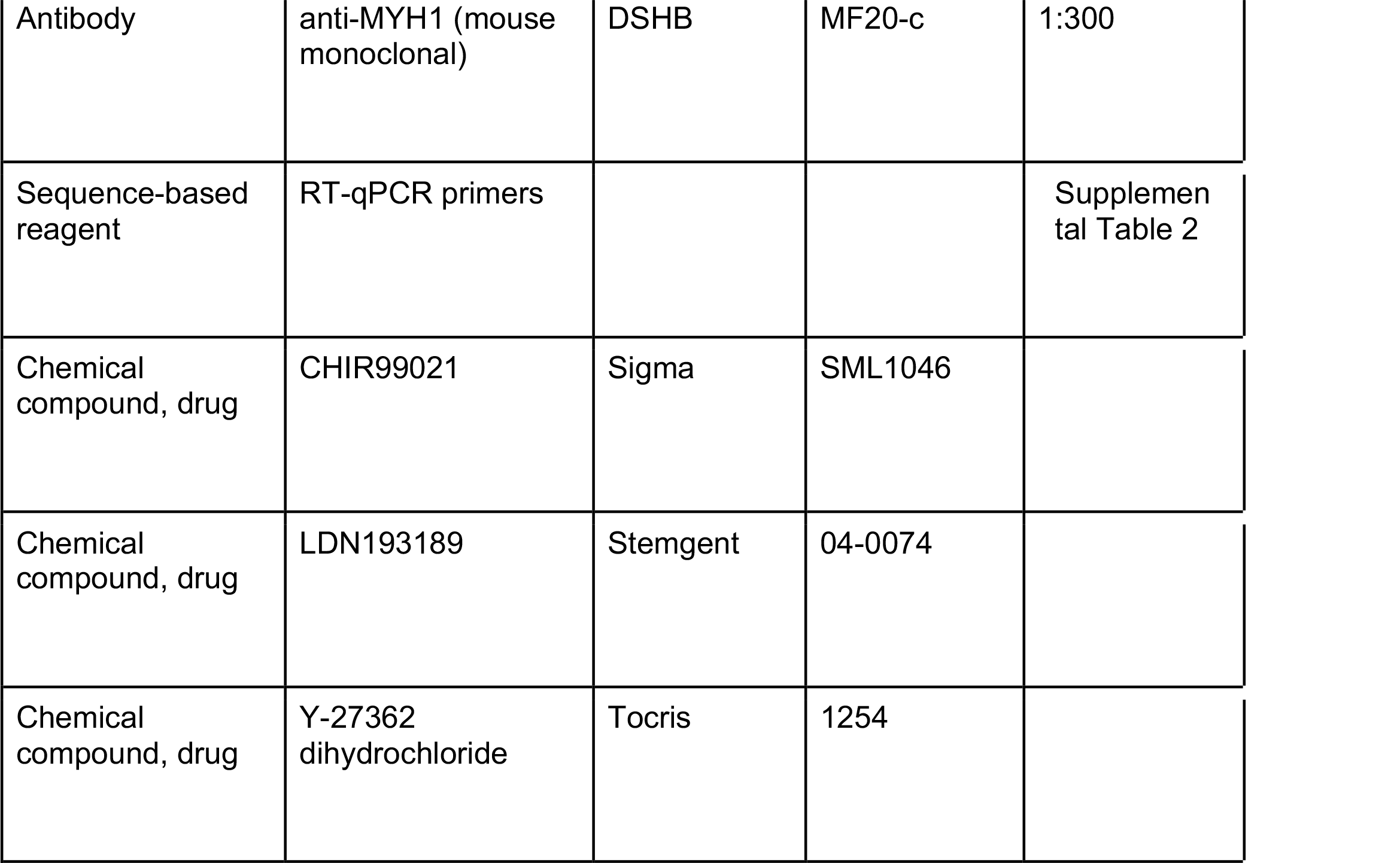

| Key Resources Table |  |  |  |  |
| --- | --- | --- | --- | --- |
| Reagent type (species) or resource | Designation | Source or reference | Identifiers | Additional information |
| Cell line (*Homo sapiens*) | NCRM1 | RUCDR Infinite Biologics | RRID:CVCL_1E71 | hiPS cell line |
| Cell line (*Homo sapiens*) | ACTB-GFP | Allen Institute | AICS-0016-184 (RRID:CVC L_JM16) | hiPS cell line |
| Cell line (*Homo sapiens*) | WTC-11 | Coriell Institute | GM25256 (RRID:CVC L_Y803) | hiPS cell line |
| Antibody | anti-CDH2/N-CADHERIN (rabbit polyclonal) | Abcam | ab18203 | 1:400 |
| Antibody | anti-PAX3 (mouse monoclonal) | DSHB | Pax3-c | 1:250 |
| Antibody | anti-SOX2 (goat polyclonal) | R&D Systems | AF2018 | 1:200 |
| Antibody | anti-T/BRACHYURY (rabbit monoclonal) | Abcam | ab209665 | 1:200 |
| Antibody | anti-TBX6 (rabbit polyclonal) | Abcam | ab38883 | 1:300 |
| Antibody | anti-TCF15 (rabbit polyclonal) | Abcam | ab204045 | 1:50 |
| Antibody | anti-MYH1 (mouse monoclonal) | DSHB | MF20-c | 1:300 |
| Sequence-based reagent | RT-qPCR primers |  |  | Supplemental Table 2 |
| Chemical compound, drug | CHIR99021 | Sigma | SML1046 |  |
| Chemical compound, drug | LDN193189 | Stemgent | 04-0074 |  |
| Chemical compound, drug | Y-27362 dihydrochloride | Tocris | 1254 |  |

### Human iPS cell culture and 3D differentiation

Human iPS cells were maintained on Matrigel-coated plates (Corning, cat no. 354277) in mTeSR1 media (Stem Cell Technologies, 85870) using maintenance procedures developed by the Allen Institute for Cell Science (https://www.coriell.org/1/AllenCellCollection). NCRM1 iPS cells were obtained from RUCDR Infinite Biologics, ACTB-GFP (cell line ID: AICS-0016 cl.184) fluorescent reporter iPS cell line was obtained from the Allen Institute for Cell Science, and the WTC-11 (GM25256) cell line was obtained from the Coriell Institute for Medical Research. All cell lines were tested for mycoplasma contamination. We verified cell line identity by immunostaining for pluripotency markers POU5F1 and SOX2.

For generation of paraxial mesoderm organoids, 500 dissociated iPS cells resuspended in mTeSR1 media containing 10 µM Y-27362 dihydrochloride (ROCKi; Tocris Bioscience, cat. no. 1254) and 0.05% poly-vinyl alcohol (PVA) were dispensed into 96-well U-bottom non-adherent suspension culture plates (Greiner Bio-One, 650185) and allowed to aggregate for 24 hours. To induce paraxial mesoderm differentiation, 24 hour old pluripotent spheroids were subjected to CL media consisting of RHB Basal media (Takara/Clontech, cat. no. Y40000), 5% KSR (Thermo Fisher Scientific, cat. no. 10828028) with 10 µM CHIR99021 (Sigma-Aldrich, cat. no. SML1046), 0.5 μM LDN193189 (Stemgent cat. no. 04-0074), and 5 µM ROCKi for the first 24 hours. Organoids were cultured in CL media without ROCKi from 24-72 hours of differentiation. On day 3 (72-120 hours), CL media was supplemented with 20 ng/ml FGF2 (PeproTech, cat. no. 450-33). On day 4, organoids were cultured in basal media only, without the addition of signaling factors.

### Human sclerotome and dermomyotome differentiation

To further differentiate Somitoids towards sclerotome fate, day 5 somite-stage organoids were treated with 5 nM of Shh agonist SAG 21k (Tocris, cat. no. 5282) and 1 µM of Wnt inhibitor C59 for 3 days as previously described (Loh et al., 2016). Organoids were subsequently differentiated towards cartilage by culturing them in the presence of 20 ng/ml BMP4 (R&D Systems, cat. no. 314-BP-010) for 6 days.

To differentiate Somitoids towards dermomyotome, day 5 somite-stage organoids were treated with CHIR99021 (3 µM), GDC0449 (150 nM) and BMP4 (50 ng/ml) for 48 hours as described previously (Loh et al., 2016; Matsuda et al., 2020).

### In vitro skeletal muscle differentiation

Day 7 Organoids differentiated towards dermomyotome fate were dissociated with Accutase, resuspended in muscle induction medium containing ROCK inhibitor Y27632 and seeded (1.5-2.5 x 10^5^ cells per well) onto Matrigel-coated 12-well plates. To induce human skeletal muscle cells, we used a N2/horse-serum containing induction medium as previously described (Matsuda et al., 2020). In brief, DMEM/F12 containing Glutamax (Thermo Fisher Scientific, Cat. No. 10565018), 1% insulin-transferrin-selenium (Thermo Fisher Scientific, Cat. No. 41400045), 1% N-2 Supplement (Thermo Fisher Scientific, Cat. No. 17502-048), 0.2 penicillin/streptomycin (Sigma-Aldrich, Cat. No. P4333-100ML), 2% horse serum (Sigma-Aldrich, Cat. No. H1270-100ML). Medium was changed every other day. Day 45 cells were fixed in 4% PFA and immunostained for Myosin heavy chain (DSHB, MF20-c, 1:300).

### Small molecule inhibitor screens

For the systematic small molecular inhibitor screen, PM organoids were generated and differentiated until day 3 (PSM) of our protocol. On day 3, media was replaced with fresh media containing combinations of small molecule inhibitors targeting the FGF, WNT, BMP and TGF-β signaling pathways at indicated concentrations. For targeting the WNT pathway, we used C59 (Tocris, cat. no. 5148), XAV939 (Tocris, cat. no. 3748) and CHIR99021 (Sigma-Aldrich, cat. no. SML1046). For inhibiting the FGF pathway we used PD173074 (Sigma-Aldrich, cat. no. P2499). For inhibiting the BMP pathway we used LDN193189 (Stemgent, cat. no. 04-0074). For inhibition of the TGF-β pathway we used A-83-01 (Tocris, cat. no. 2939). Media was changed daily. We analyzed 3 replicates per condition in the primary screen, and 5 replicates per condition in the secondary screen.

### Immunostaining

For organoid whole-mount immunostaining, organoids were collected in cold PBS and fixed in 4% paraformaldehyde for 1-2 hours depending on size/stage. Organoids were washed in PBS and PBSFT (PBS, 0.2% Triton X-100, 10% FBS), and blocked in PBSFT+3% normal donkey serum. Primary antibody incubation was performed in the blocking buffer overnight at 4°C on a rocking platform. After extensive washes in PBSFT, secondary antibody incubation (1:500, all secondary antibodies were raised in donkey) was performed overnight in PBSFT. Organoids were washed first in PBSFT and, for the final washes, were transferred to PBT (PBS, 0.2% Triton X-100, 0.2% BSA), followed by 50% glycerol in PBT and 70% glycerol in PBT prior to mounting. Hoechst (1:2000) was added to the last PBSFT wash. A list of primary antibodies is provided in Table S1.

### Confocal and time-lapse microscopy

All whole-mount immunostaining images were collected with a Nikon A1R point scanning confocal with spectral detection and resonant scanner on a Nikon Ti-E inverted microscope equipped with a Plan Apo VC 20x objective (NA 0.75). Alexa-488, Alexa-594, Alexa-647 fluorophores coupled to secondary antibodies were excited with the 488 nm, 561 nm, and 647 nm laser lines from a Spectral Applied Research LMM-5 laser merge module with solid state lasers (selected with an AOTF) and collected with a 405/488/561/647 quad dichroic mirror (Chroma). For time-lapse experiments, images were acquired with a Yokagawa CSU-X1 spinning disk confocal on a Nikon Ti inverted microscope equipped with a Plan Apo 20x objective (NA 0.75) and a Hamamatsu Flash4.0 V3 sCMOS camera. Samples were grown on 6-well glass-bottom multiwell plates with No. 1.5 glass (Cellvis, cat. no. P06-1.5H-N) and mounted in a OkoLab 37°C, 5% CO2 cage microscope incubator warmed to 37°C. Images were collected every 15 min, using an exposure time of 800 ms. At each time-point, 30 z-series optical sections were collected with a step-size of 2 µm. Multiple stage positions were collected using a Prior Proscan II motorized stage. Z-series are displayed as maximum z-projections, and gamma, brightness, and contrast were adjusted (identically for compared image sets) using Fiji/ImageJ (Schindelin et al., 2012; https://imagej.net/Fiji).

### Automated image segmentation and analysis

Automated image analysis, including background de-noising, segmentation and feature extraction was done using ImageJ/Fiji macro language run in batch mode to process the entire screen data set. First, binary masks were generated from the Hoechst (nuclear stain) channel by de-noising the image (Gaussian Blur, sigma=5) followed by applying Li’s Minimum Cross Entropy thresholding method (Li and Tam, 1998) and refining binary masks through several rounds of erosion/dilation steps. Next, binary masks were converted to selections and added to the Region of Interest (ROI) Manager. Finally, ROIs were used to perform diameter measurements of organoids. For Pax3 measurements, Hoechst and Pax3 channels were first denoised using a Gaussian blur filter (sigma=10) and then used to create a normalized Pax3 image by dividing the Pax3 channel with the Hoechst channel. Next, ROIs based on Hoechst binary masks were applied to the Pax3 normalized image to extract fluorescence intensity measurements for each z-slice. Finally, mean Pax3 intensity values for each organoid were calculated and compared.

### Quantification of per organoid somite numbers for secondary screen

For the primary and secondary screens, images were acquired on a Nikon A1R point scanning confocal microscope. For each organoid, 66 z-series optical sections were collected with a step-size of 2 µm. Somite quantification for the secondary screen was done by blinded manual scoring, considering the following criteria:

1. Nuclear expression of somitic marker PAX3.
2. Accumulation of NCAD around a central cavity.
3. Radial arrangement of PAX3+ columnar cells around the central cavity (rosette-like structure).

### Quantification of organoid and human somite sizes

Carnegie stage 9, 10 and 11 human embryonic somite data was obtained from the Virtual Human Embryo Project (https://www.ehd.org/virtual-human-embryo/). Somite sizes of human embryos were measured using the Ruler Tool on the Virtual Human Embryo website along the medio-lateral and dorso-ventral axis of the embryo. The slice with the largest diameter of each somite was used for measurements. Somite sizes of day 5 organoids were measured along the X and Y axes of the image since, unlike in the embryo, they do not exhibit morphological anisotropies. Somite areas were approximated by using the two diameter measurements from each somite to calculate the area of the resulting rectangle.

### RNA extraction, reverse transcription and qPCR

Organoids were harvested in Trizol (Life Technologies cat. no. 15596-018), followed by precipitation with Chloroform and Ethanol and transfer onto Purelink RNA Micro Kit columns (Thermo Fisher cat. no. 12183016) according to manufacturer’s protocol, including on-column DNase treatment. A volume of 22 µl RNase-free water was used for elution and RNA concentration was measured with a Qubit Fluorometer. Typically, between 0.2-1 µg of RNA was reverse transcribed using Superscript III First Strand Synthesis kit (Life Technologies cat. no. 18080-051) and oligo-dT primers to generate cDNA libraries.

For real time quantitative PCR, cDNA was diluted 1:30-1:50 in water and qPCR was performed using the iTaq Universal SYBR Green kit (Bio-Rad cat. no. 1725124). Each gene-specific primer and sample mix was run in triple replicates. Each 10 µl reaction contained 5 µl 2X SYBR Green Master Mix, 0.4 µl of 10 µM primer stock (1:1 mix of forward and reverse primers), and 4.6 µl of diluted cDNA. qPCR plates were run on a Roche LightCycler 480 Real-Time PCR system with the following cycling parameters: initial denaturation step (95°C for 1 minute), 40 cycles of amplification and SYBR green signal detection (denaturation at 95°C for 5 seconds, annealing/extension and plate read at 60°C for 40 seconds), followed by final rounds of gradient annealing from 65°C to 95°C to generate dissociation curves. Primer sequences are listed in Table S2. All unpublished primers were validated by checking for specificity (single peak in melting curve) and linearity of amplification (serially diluted cDNA samples). For relative gene expression analysis, the ΔΔCt method was implemented using the R package ‘pcr’ (https://cran.r-project.org/web/packages/pcr/). PP1A was used as the housekeeping gene in all cases. Target gene expression is expressed as fold change relative to undifferentiated iPS cells.

### Preparation of single-cell suspensions for scRNA-seq

Cell dissociation protocols were optimized to achieve single cell suspensions with >90% viable cells and low number of doublets. Organoids collected at day 1, 2, 3, and 5 of our differentiation protocol were pooled in pre-warmed PBS, transferred to pre-warmed accutase and incubated for 5-7 min at 37°C. For Day 1 organoid cell suspension, 30 organoids were pooled. For Day 2 organoid cell suspension, 15 organoids were pooled. For Day 3 cell suspension, 8 organoids were pooled. For Day 5 cell suspension, 5 organoids were pooled. Organoids were briefly rinsed in PBS, then transferred to 500 µl PBS/0.05% BSA and carefully triturated to generate a single-cell suspension. Cell suspension was run through a cell strainer (Falcon, cat. no. 352235) and transferred to a 1.5 ml tube. Cells were spun down at 250g for 3 min at 4°C. Cell pellet was resuspended in 25 µl PBS/0.05% BSA, cell concentration and viability was measured using an automated cell counter, and cell suspension was further diluted as appropriate to reach the optimal range for 10x (700-1200 cells per µl). Cells were subjected to single-cell RNA sequencing (10x Genomics, Chromium Single Cell 3’ v3) aiming for the following target cell numbers: Day 1, 3,000 cells; Day 2, 4,000 cells; Day 3, 5,000 cells; Day 5, 6,000 cells. Estimated actual cell numbers collected were: Day 1, 2,930 cells; Day 2, 4,977; Day 3, 5,968 cells; Day 5, 4,841 cells. Single-cell libraries were generated using standard protocols. Libraries were sequenced together on a NovaSeq 6000 system resulting in 800 million reads.

### Analysis of scRNA-seq data

Statistics and plots were generated using R version 4.0.2 “Taking Off Again” and Seurat version 3.0 (Stuart et al., 2019).

### QC analysis / processing of scRNA-seq data

Cell Ranger pipeline (10x Genomics, Version 4.0.0) was used to de-multiplex the raw base call files, generate FASTQ files, perform the alignment against the human reference genome (GRCh38 1.2.0), and generate the count matrices.

For the initial QC, we determined the following thresholds for filtering out low-quality cells: UMI counts less than 500, gene counts less than 200, mitochondrial fraction above 0.2 and a complexity score of less than 0.8 (calculated as log10(genes)/log10(UMIs)).

### Low-dimensional embedding and clustering

After QC filtering, we normalized our dataset using the sctransform (Hafemeister and Satija, 2019) framework, which is part of the Seurat package. To regress out confounding variation in our dataset, we performed cell cycle scoring and determined mitochondrial mapping percentage using standard workflows. Next, we performed PCA and determined the K-nearest neighbor graph using the first 40 principle components. We then applied the Leiden clustering algorithm using a parameter range from 0.1-1.0 to determine the best resolution/number of clusters, which reflected biological differences (FindClusters, resolution=0.1-1.0). Clusters were visualized on a UMAP embedding (RunUMAP, dims = 1:40). To determine optimal resolution for clustering and assign cell types for each cluster, we visualized sets of known marker genes for each predicted cell type on UMAP plots. Prior to marker gene identification and final assignments of cluster identities, we also checked additional quality control metrics (UMI count, gene count, mitochondrial gene ratio) to exclude low-quality clusters from downstream analyses. Through iterative analysis we determined Leiden clustering with resolution = 0.8, resulting in 22 clusters, to best capture biological variation of the dataset. Using a combination of quality control metrics and unbiased marker gene identification for each cluster (see below), we excluded seven smaller low-quality clusters (as determined by QC metrics and/or expression of stress signature genes) from further downstream analysis (15 clusters after filtering).

### Identification of differentially expressed genes

Marker genes for every cluster were identified by a two-sided Wilcoxon rank-sum test comparing cells from each cluster to all other cells in the combined dataset. Genes were considered differentially expressed if the log2 fold-change average expression in the cluster is equal or greater 0.25 relative to the average expression in all other clusters combined, and the adjusted P-value <0.05. Multiple comparison correction was performed using the Bonferroni method. Identified marker genes for the top 20 differentially expressed transcripts are listed in Fig 4-Supplemental Fig 4A. The full list of differentially expressed genes, ranked by adjusted P-values and associated fold-changes are provided in Supplemental Table 1.

---

**Figure 1-Supplemental Figure 1.**
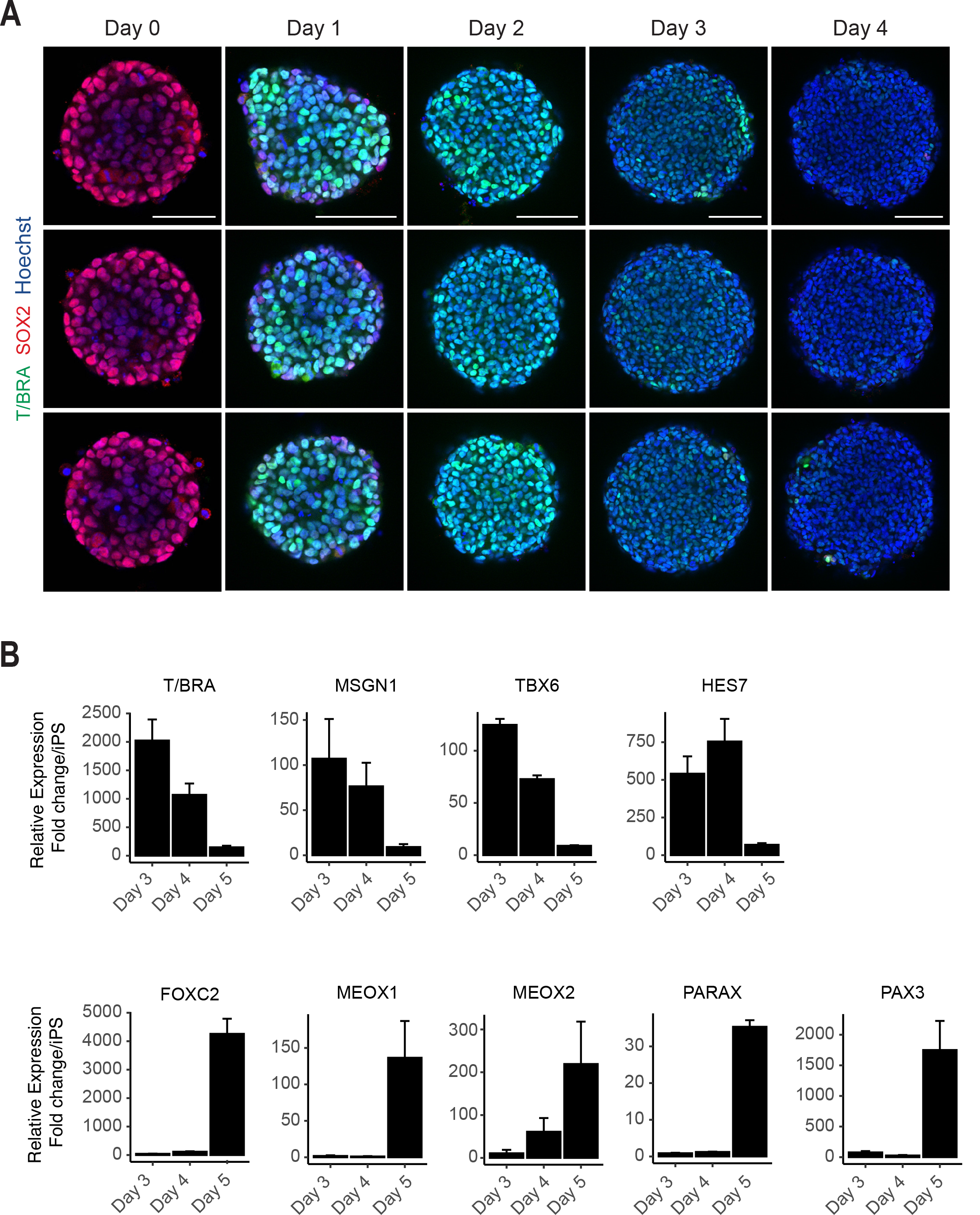
Additional immunofluorescent and qPCR data. (A) Immunofluorescent images of PM organoid differentiation from cavity-forming spheroids. Single optical z-sections are shown for each organoid to illustrate the central cavity formed 24 hours after aggregation (Day 0). For each day of differentiation, three representative replicates are shown. Representative images are shown from 3 organoid replicates. NCRM1 hiPS cells were used to generate organoids. (B) qRT-PCR analysis of PSM and Somite markers reveals PSM-to-somite transition from day 4 to day 5. Relative gene expression levels are shown as fold-change relative to undifferentiated iPS cells (see Methods).

**Figure 1-Supplemental Figure 2.**
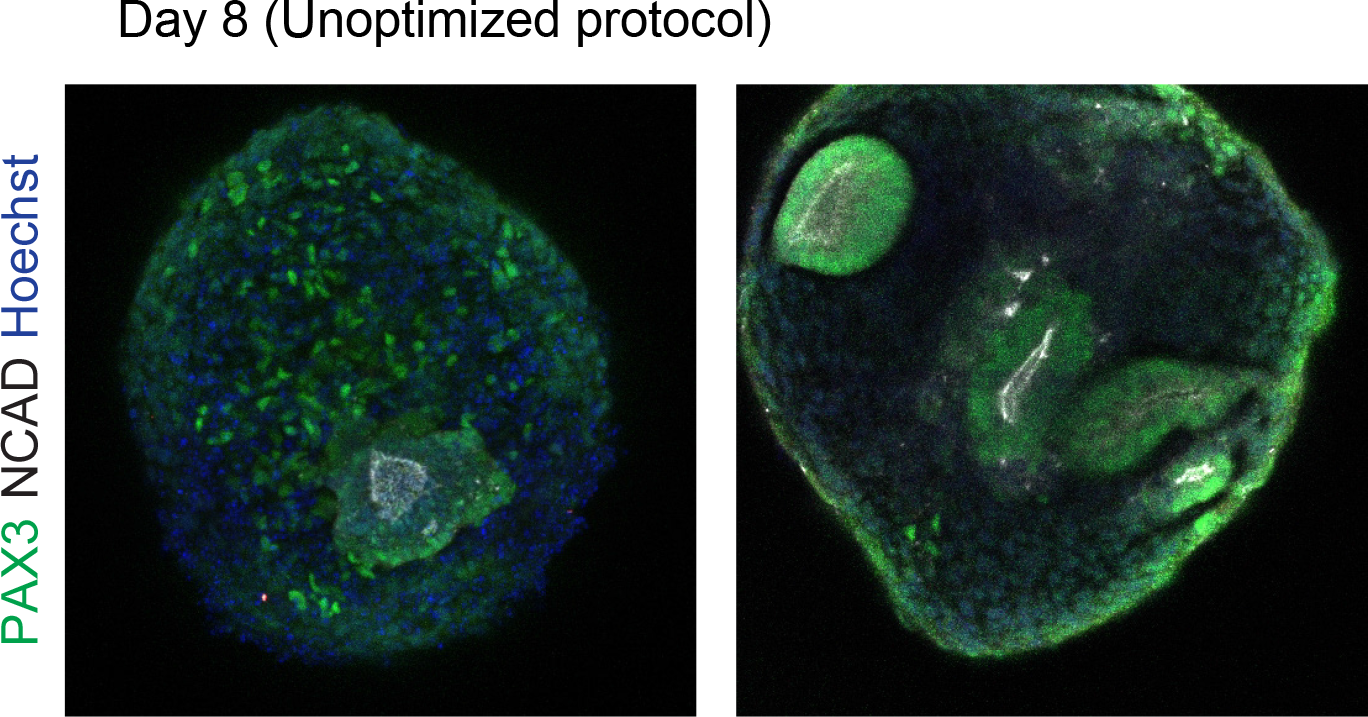
Organoids generated using an unoptimized protocol exhibit heterogeneous activation of somite marker genes (PAX3 and NCAD) and a low number of rosette structures. The rosette structures do not match the expected size of *in vivo* human somites. In addition, organoids generated using the unoptimized protocol showed a delayed onset of PAX3 expression and the rosette structures did not form until day 8.

**Figure 2-Supplemental Figure 1.**
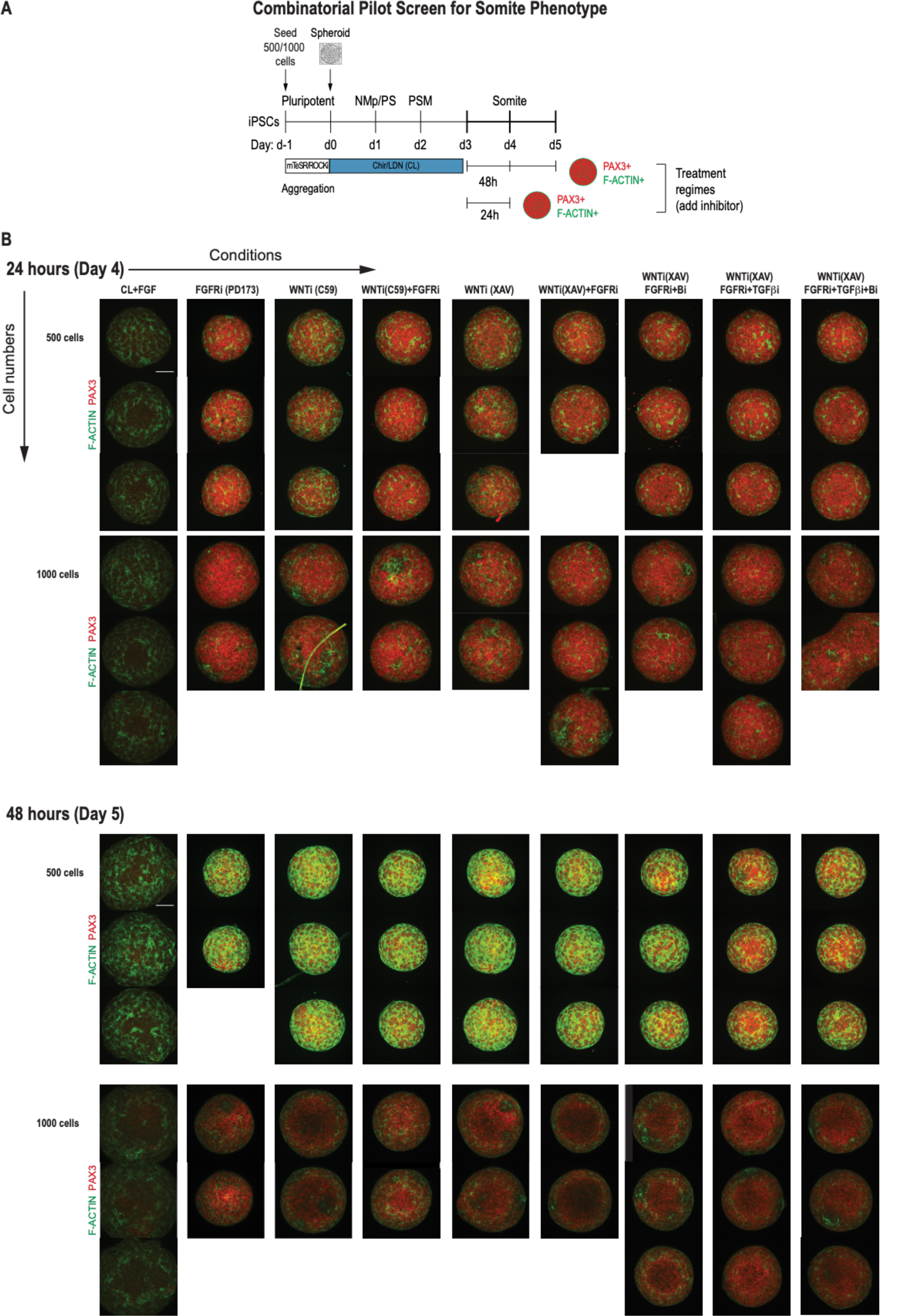
Replicate data of pilot screen for somite phenotype in human PM organoids. (A) Schematic overview of systematic screen in PM organoids. PM-stage organoids were treated on day 3 for 24 hours and 48 hours, respectively, with signaling agonists/antagonists as indicated. Control organoids were maintained in CL media with FGF added. (B) Complete immunofluorescent data set showing all replicates of day 4 and day 5 organoids after treatment immunostained for somite marker PAX3 and F-ACTIN to visualize rosette-like somite structures. Confocal images are shown as maximum intensity z-projections. Scale bar represents 100 µm. Small molecule inhibitors used are indicated in brackets. FGFRi: FGF receptor inhibitor (PD173074). WNTi: Wnt inhibitor (C59 or XAV939). Bi: BMP inhibitor (LDN). TGF-βi: TGF-β inhibitor (A-83-01).

**Figure 2-Supplemental Figure 2.**
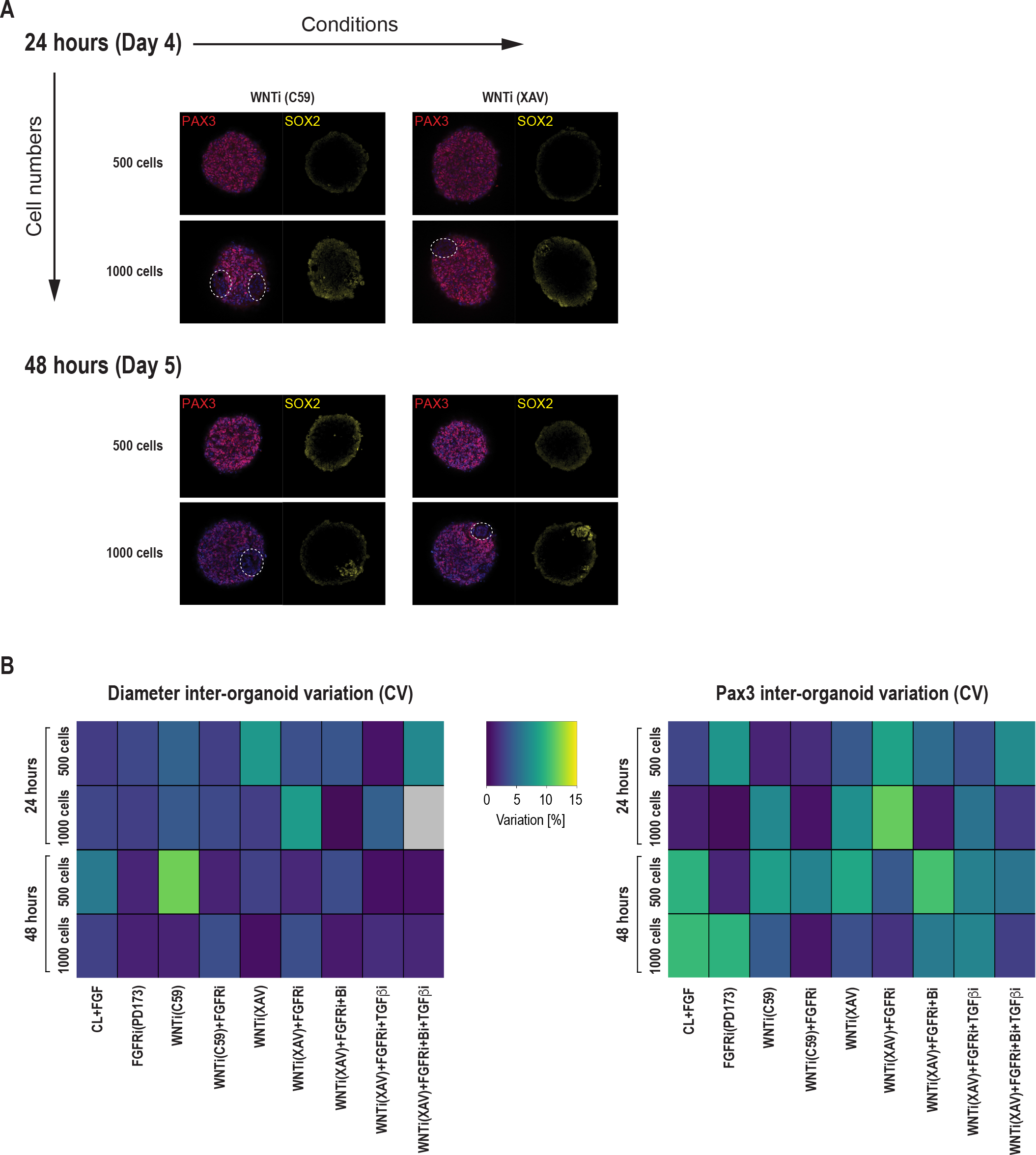
Additional immunofluorescent data and inter-organoid phenotypic variance of primary somite phenotype screen in Somitoids. (A) Comparison of organoids made from 500 and 1000 cells, respectively, immunostained for PAX3 (somite fate) and SOX2 (neural fate). Organoids made from 1000 cells contain patches of PAX3-negative/SOX2-positive cells (marked by dashed contours), indicating a higher degree of heterogeneity of somite fate induction. Representative images are shown from 3 organoid replicates. (B) Heatmaps of diameter and PAX3 inter-organoid variation for each treatment, expressed as coefficient of variation (CV, in percentage). CV is calculated across the 2 or 3 organoid replicates of each condition as (standard deviation/mean)*100.

**Figure 2-Supplemental Figure 3.**
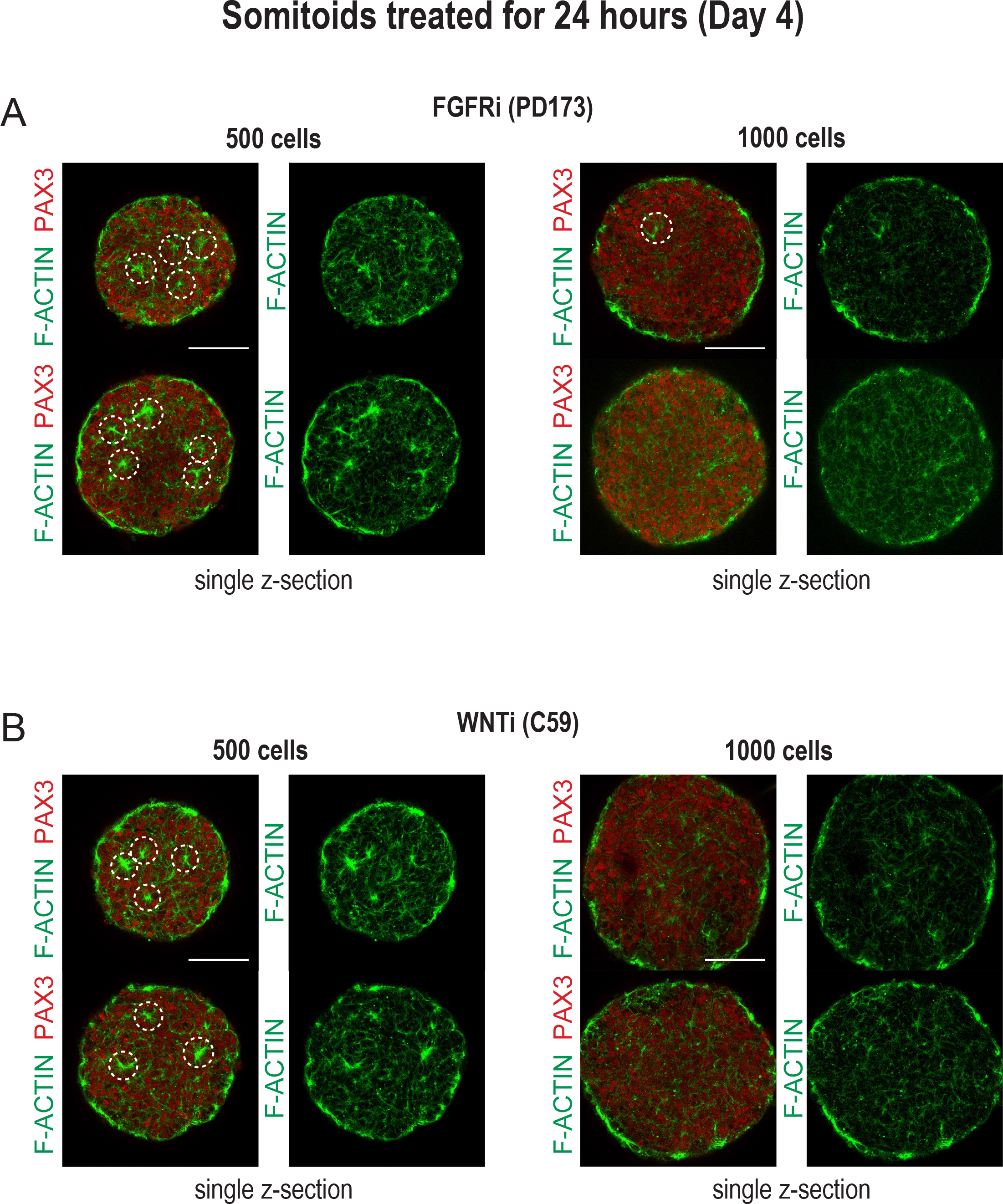
Organoids made from 500 cells more reproducibly generate somite-like structures compared to organoids made from 1000 cells. Representative single z-sections of organoids treated for 24 hours with inhibitors as indicated. Organoids were stained for PAX3 and F-ACTIN to visualize somite-like structures. Scale bar represents 100 µm. (A) Organoids made from 500 and 1000 cells, respectively, were treated with FGFR inhibitor PD173 for 24 hours on day 3 of the differentiation protocol. Two representative organoids are shown for each cell number. Organoids made from 500 cells consistently show more somite-like structures compared to organoids made from 1000 cells. (B) Organoids made from 500 and 1000 cells, respectively, were treated with WNT inhibitor C59 for 24 hours on day 3 of the differentiation protocol. Two representative organoids are shown for each cell number. Organoids made from 500 cells show more uniform expression of PAX3 and a higher number of somite-like structures compared to organoids made from 1000 cells.

**Figure 3-Supplemental Figure 1.**
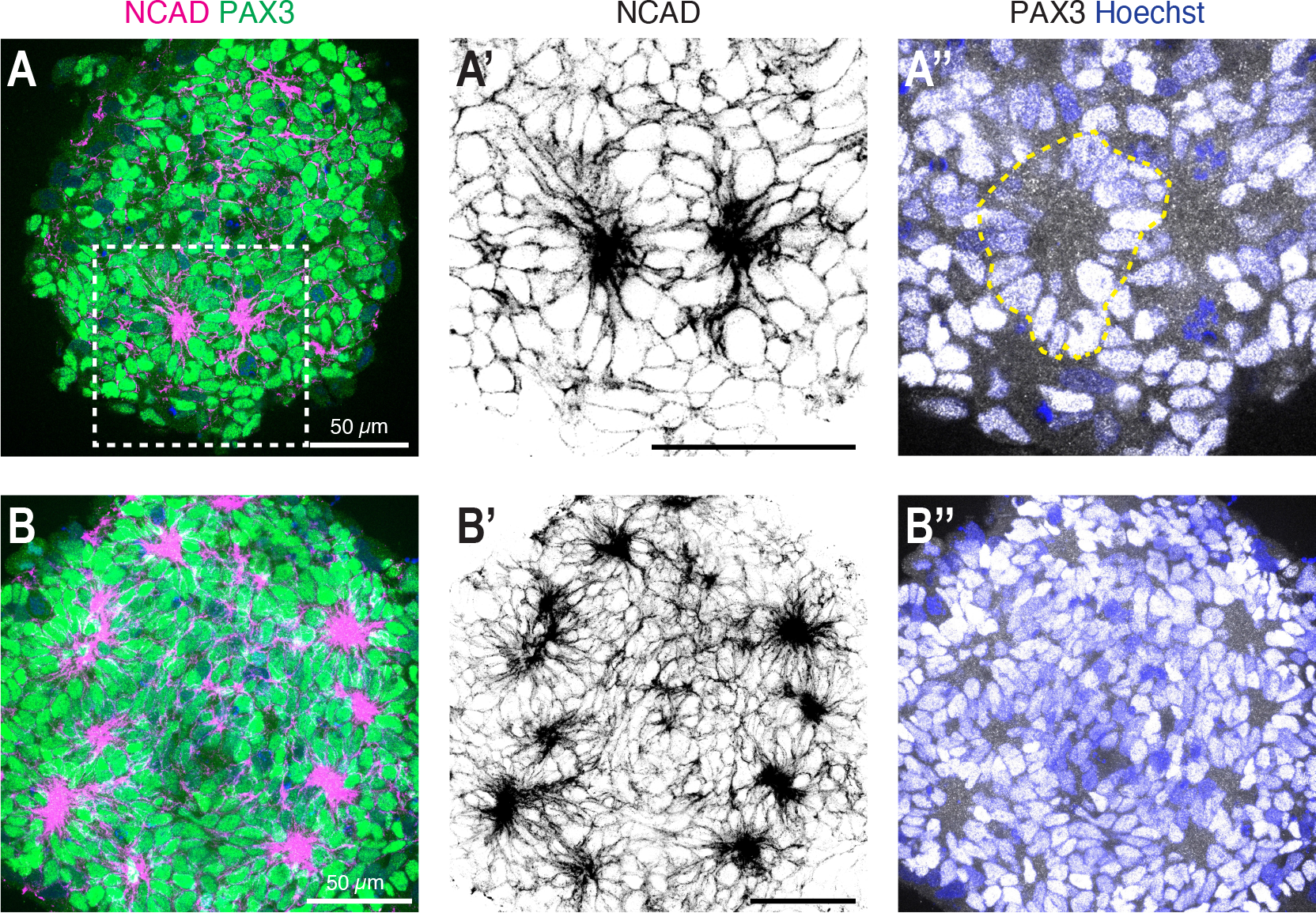
High-resolution imaging of somite-like structures in day 5 organoids. (A) Single z-section of a day 5 organoid treated with CL+FGF2 on day 3 to day 4, followed by culture in basal media for 24 hours (optimized protocol). Somites were visualized by immunostaining using somite markers PAX3 and NCAD. (A’) shows an enlargement of the dashed square indicated in (A) with two somites visible based on NCAD expression. (A’’) Merged image of the PAX3 and Hoechst (nuclear stain) channels with a dashed outline marking one of the somites. (B) Z-projection of a day 5 organoid treated with CL+FGF2 on day 3 to day 4, followed by culture in basal media for 24 hours (optimized protocol). 5 consecutive z-sections (1 µm step size) were used to generate the projection. Somites were visualized by immunostaining using somite markers PAX3 and NCAD. Scale bars indicate 50 µm.

**Figure 3-Supplemental Figure 2.**
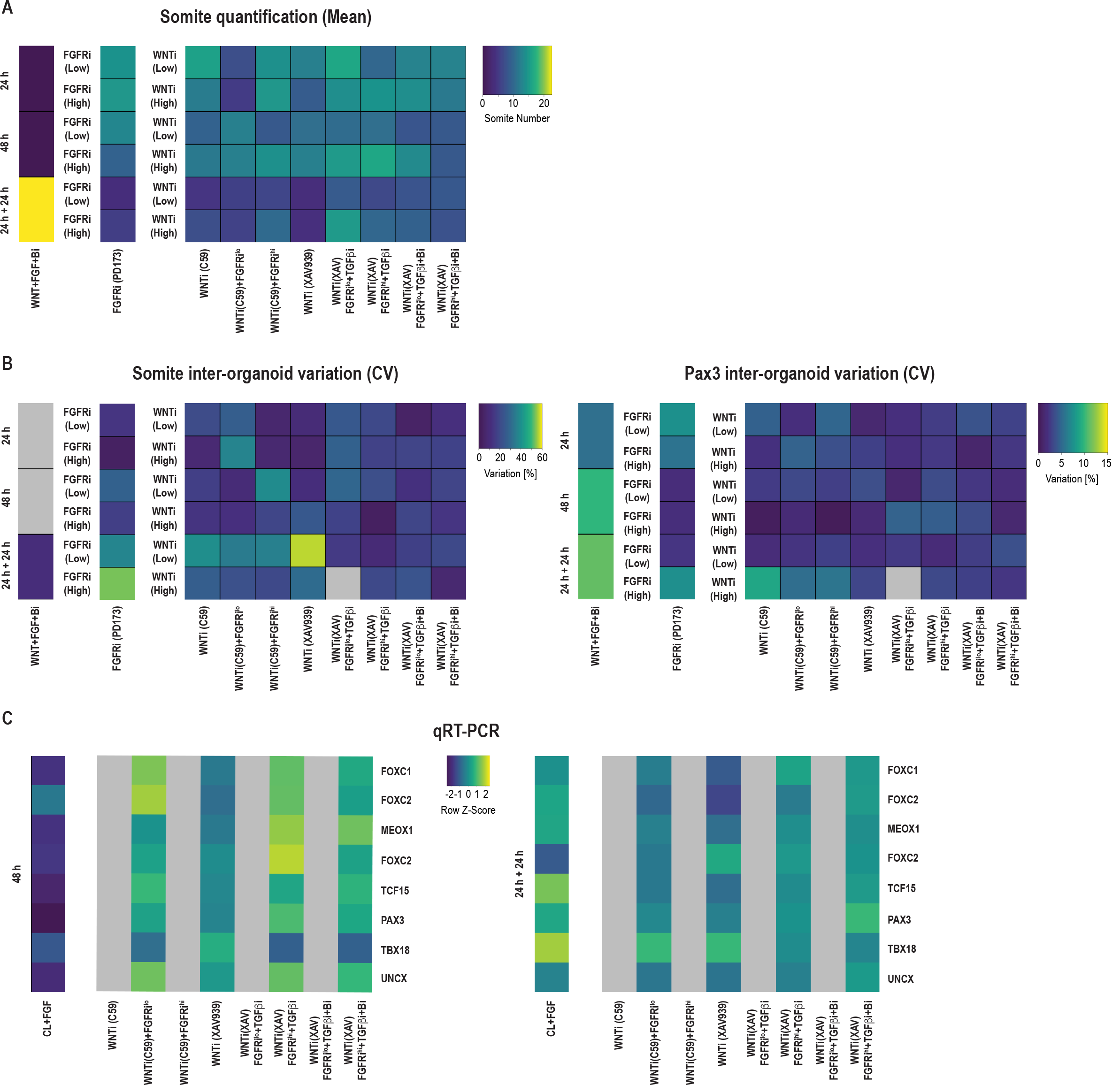
Additional qPCR data and inter-organoid phenotypic variance of secondary somite phenotype screen in somitoids. (A) Quantification of the number of somite-like structures for each condition. Somite numbers are shown as calculated means based on quantifying somites for 2-5 organoids per condition. (B) Heatmaps of Somite count and PAX3 inter-organoid variation for each treatment, expressed as coefficient of variation (CV, in percentage). CV is calculated across the 3-5 replicates of each condition as (standard deviation/mean)*100. (C) qRT-PCR analysis of somite marker genes for select treatment conditions from the secondary somite phenotype screen. Expression of somite marker genes correlates well with Pax3 immunostainings from the secondary screen (see Fig 3B). Relative gene expression levels are shown as Z-scores, expressed as fold-change relative to undifferentiated iPS cells (see Methods). Source data is available in Fig 3-Supplemental Fig 2-Source Data 1.

**Figure 3-Supplemental Figure 3.**
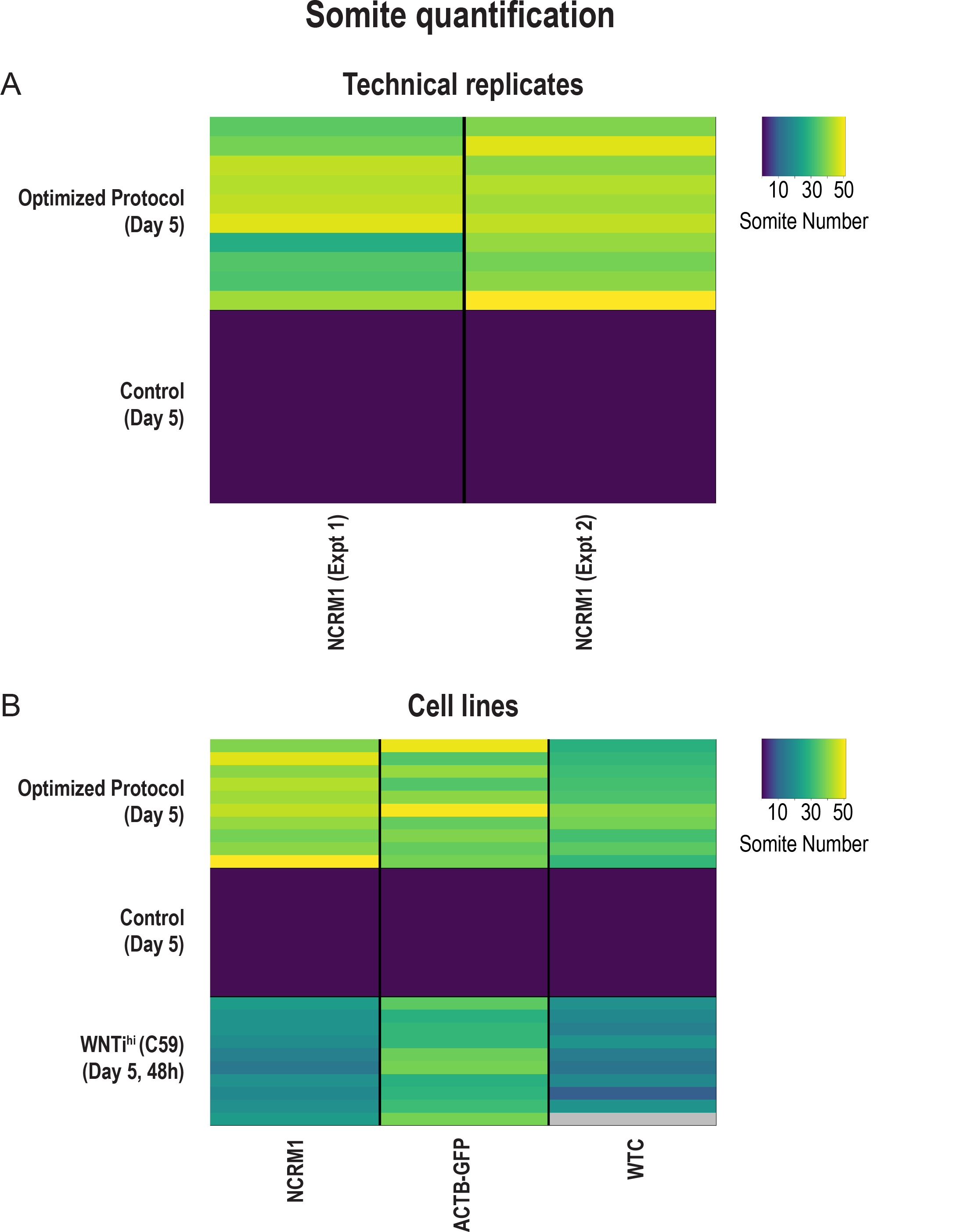
The optimized Somitoid protocol is reproducible across experiments and different cell lines. Quantification of the number of somite-like structures for each condition. Each row represents one organoid replicate. 10 organoids are shown per condition. Optimized protocol corresponds to treatment with CL+FGF2 for 24 hours followed by culture in basal media for 24 hours. Control organoids were treated with CL+FGF2 for 48 hours. (A) Organoids treated with the optimized protocol exhibit a robust and reproducible somite phenotype across two independent experiments (Experiment 1, average somite number = 39+/-8 (mean+/-std); Experiment 2, average somite number = 42+/-4; p-val = 0.16). Source data is available in Fig 3-Supplemental Fig 3-Source Data 1. (B) Applying the optimized protocol to multiple cell lines confirms that the observed somite phenotype is reproducible across different cell lines. Two cell lines (ACTB-GFP and WTC) were characterized in addition to our screening cell line (NCRM1). NCRM1, average somite number = 43+/-4 (mean+/-std); ACTB-GFP, average somite number = 40+/-6; WTC, average somite number = 33+/-4. One additional experimental condition (C59^hi^, 48h), which scored amongst the best conditions in our secondary screen, was also tested and compared across cell lines. Although the C59 treatment (a WNT inhibitor) resulted in a higher number of somites in the ACTB-GFP cell line (average somite number = 32+/-4 (mean+/-std)) compared with the other cell lines (NCRM1, average somite number = 20+/-3; WTC, average somite number = 17+/-4), the optimized protocol resulted in the highest number of somites in all cell lines. Source data is available in Fig 3-Supplemental Fig 3-Source Data 2.

**Figure 4-Supplemental Figure 1.**
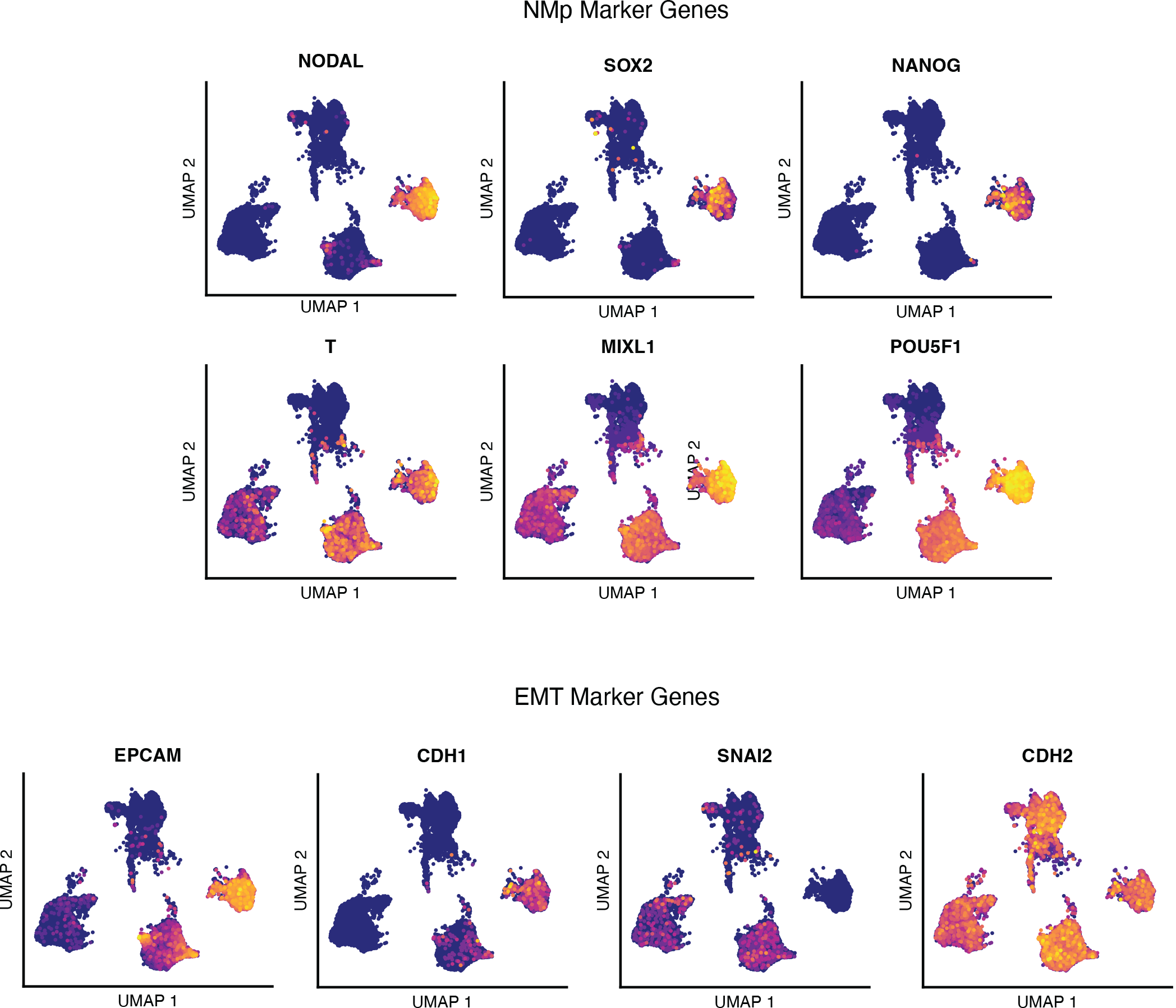
Single-cell RNA sequencing analysis of differentiating human PM organoids, NMp marker genes. UMAP plots of single-cell transcriptomes of differentiating human PM organoids overlaid with normalized transcript counts of selected marker genes. NMp, neuro-mesodermal progenitor, EMT, epithelial-to-mesenchymal transition.

**Figure 4-Supplemental Figure 2.**
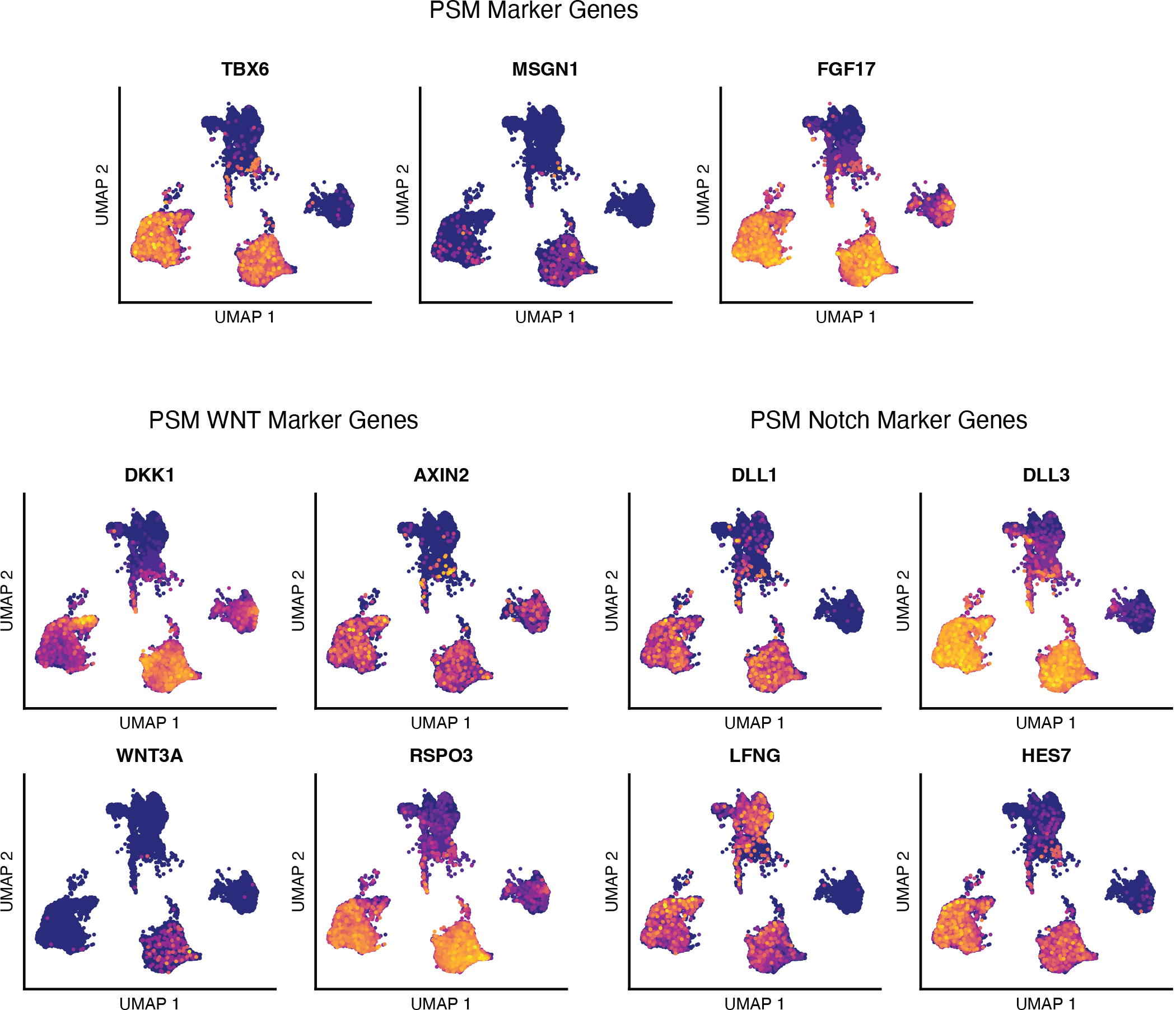
Single-cell RNA sequencing analysis of differentiating human PM organoids, PSM marker genes. UMAP plots of single-cell transcriptomes of differentiating human PM organoids overlaid with normalized transcript counts of selected marker genes. PSM, pre-somitic mesoderm.

**Figure 4-Supplemental Figure 3.**
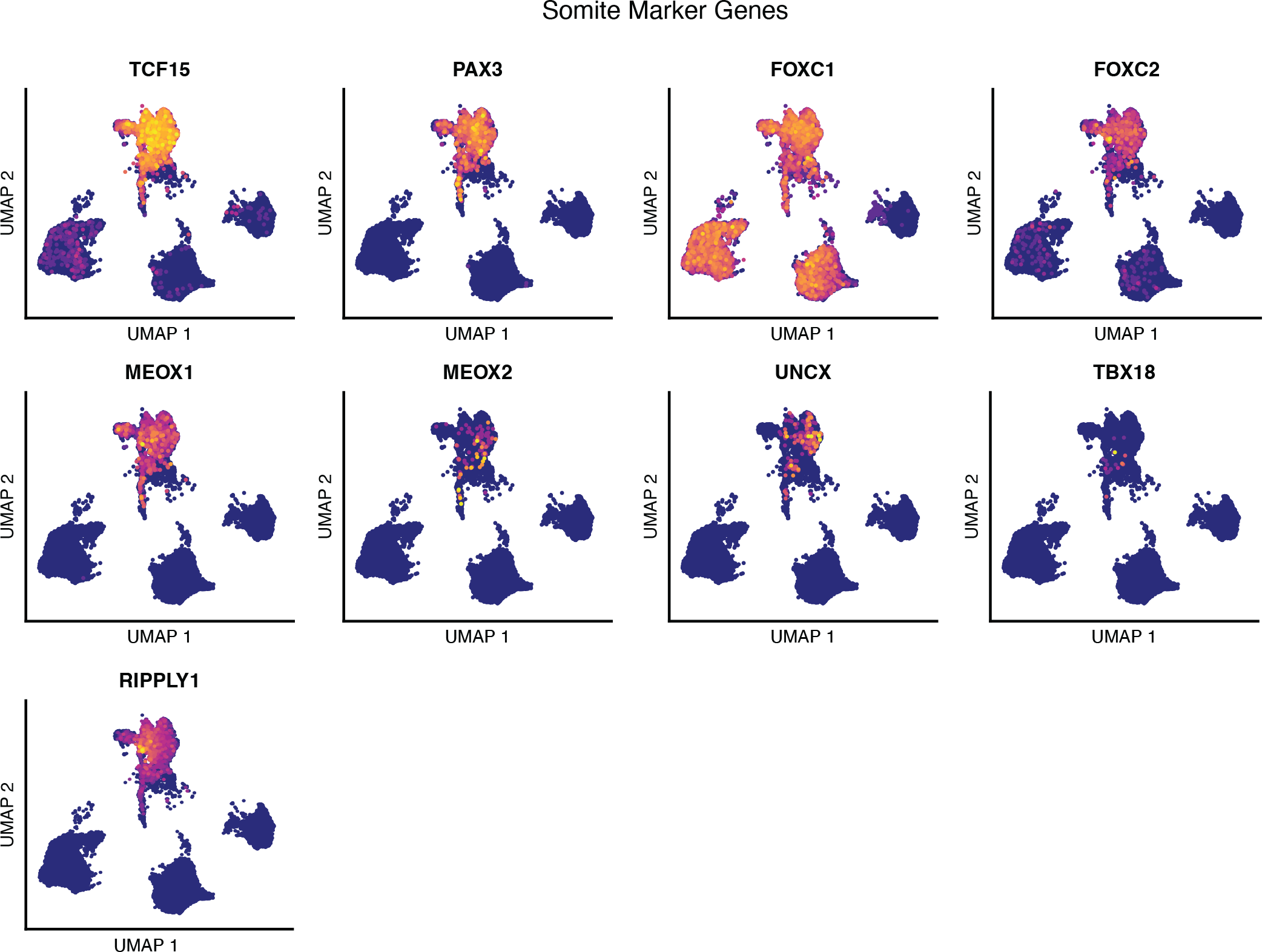
Single-cell RNA sequencing analysis of differentiating human PM organoids, Somite marker genes. UMAP plots of single-cell transcriptomes of differentiating human PM organoids overlaid with normalized transcript counts of selected marker genes.

**Figure 4-Supplemental Figure 4.**
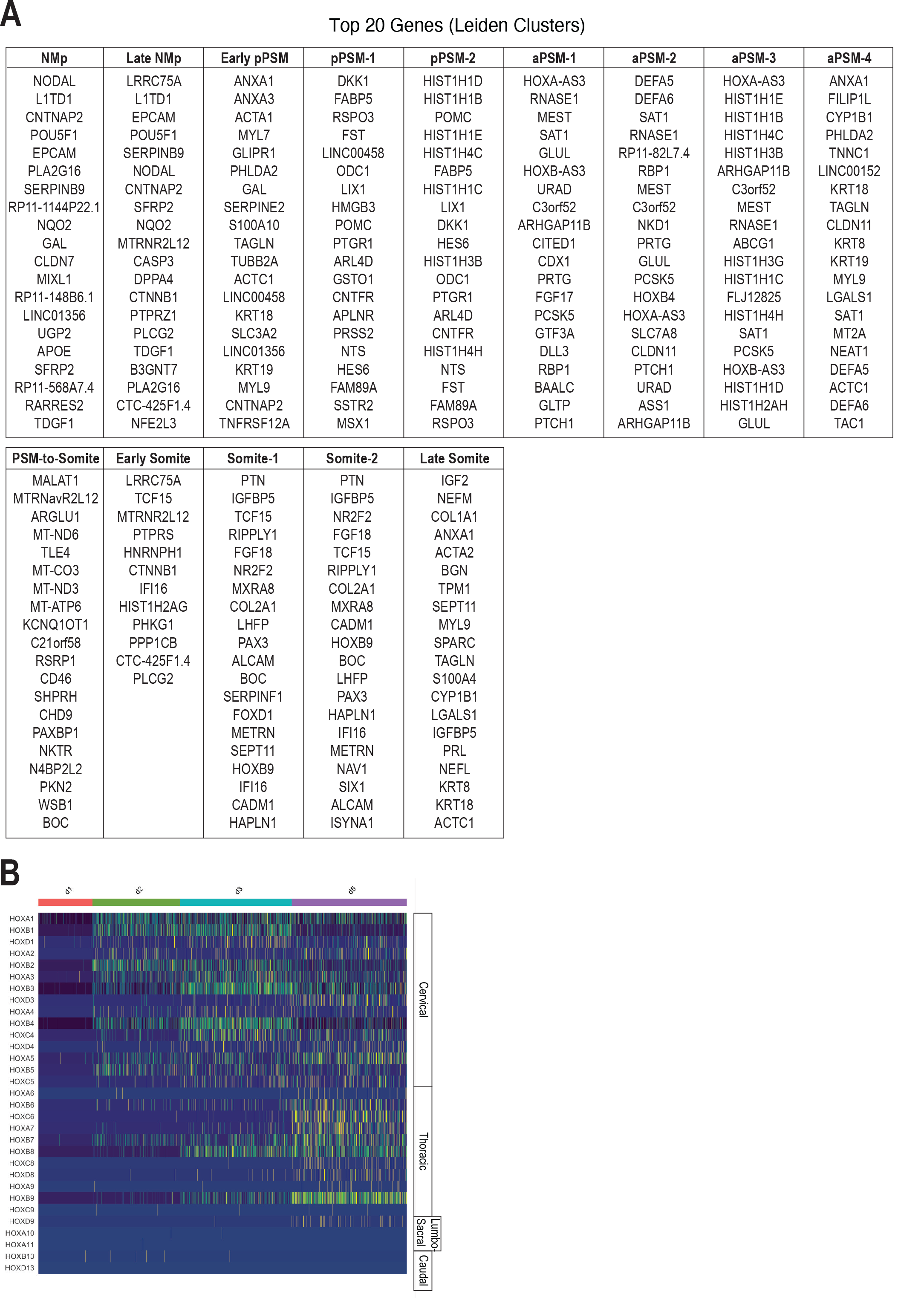
Single-cell RNA sequencing analysis of differentiating human PM organoids, cluster-based marker gene identification and HOX gene analysis. (A) Top 20 positively enriched genes for identified Leiden clusters relative to all other clusters. Genes were identified by a two-sided Wilcoxon rank-sum test and are ranked by adjusted P values based on Bonferroni correction. See Supplemental Table 1 for a complete list of identified marker genes for each cluster, adjusted P-values and fold-change values. (B) Heat map of single-cell HOX gene expression levels. Cells are grouped by collection time point. Hox genes are ordered by position, with anatomical positions of HOX paralogues indicated on the right.

**Figure 4-Supplemental Figure 5.**
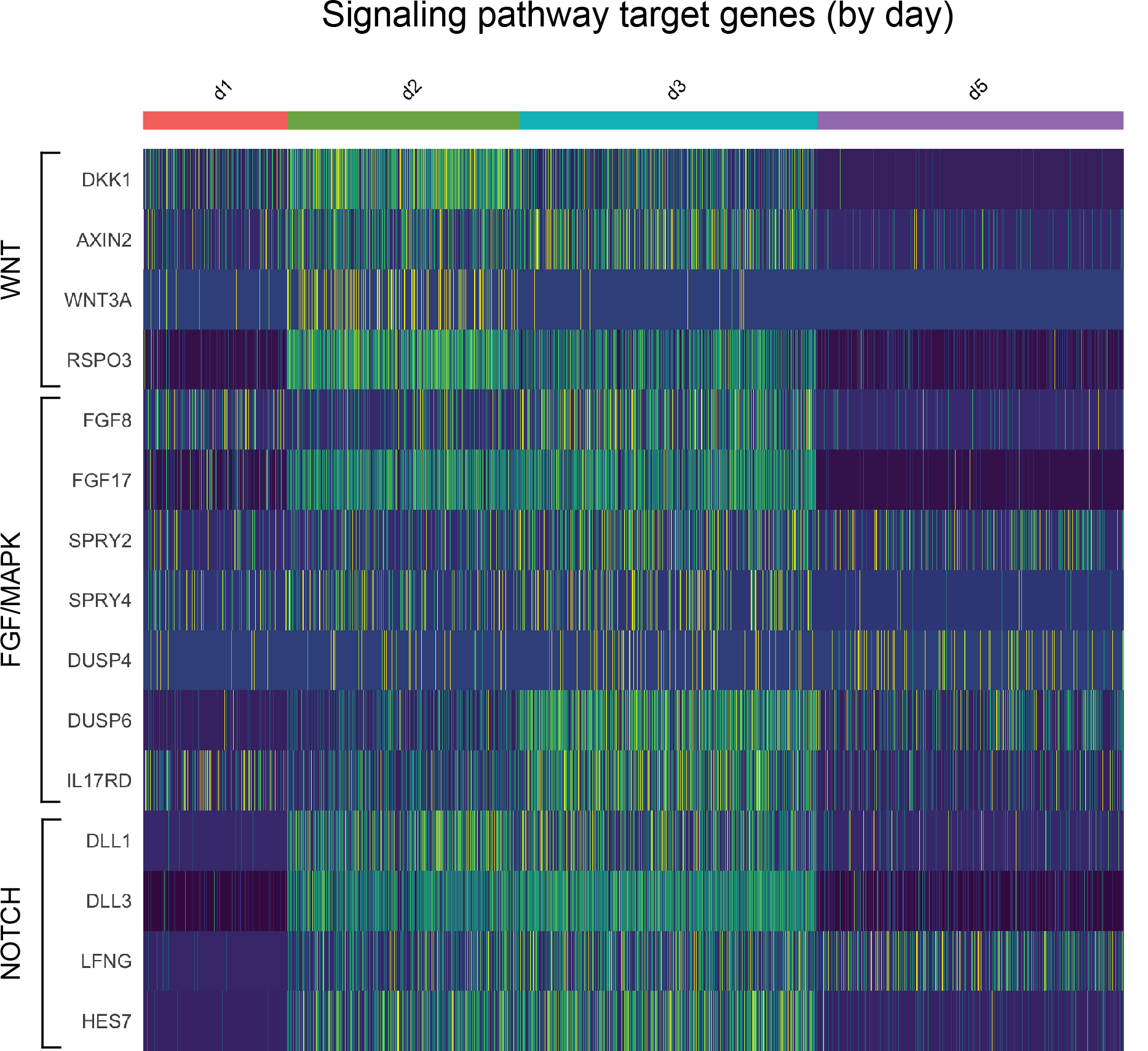
Single-cell RNA sequencing analysis of differentiating Somitoids reveals downregulation of WNT, FGF, and NOTCH target genes in day 5 somite-like cells. Heat map of single-cell signaling target gene expression levels. Cells are grouped by collection time point. Genes are grouped based on their signaling pathway, indicated on the left. Day 5 somitic cells autonomously downregulate many WNT, FGF, and NOTCH signaling target genes.

**Figure 5-Supplemental Figure 1.**
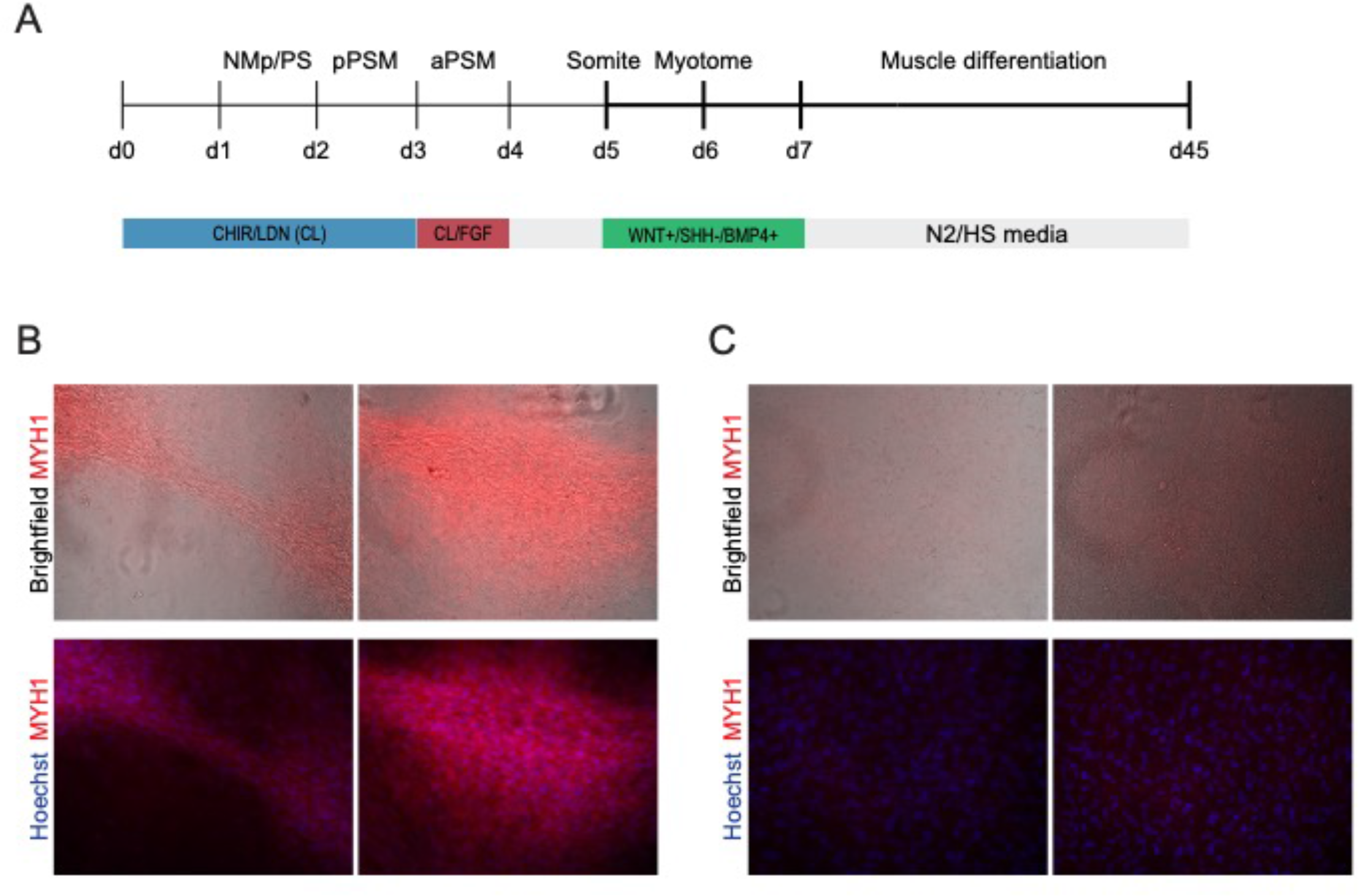
Differentiation of Somitoid-derived cells towards skeletal muscle. (A) Day 5 somite-stage organoids were further differentiated towards skeletal muscle. Organoids were first exposed to signaling modulators of WNT, SHH, and BMP4 for 48 hours to differentiate them to dermomyotome. Next, organoids were dissociated, and further differentiated as monolayer cultures on Matrigel in muscle differentiation medium. Day 5 control organoids were cultured for an additional 48 hours in basal media (day 5-7) and not pulse-treated with WNT+/SHH-/BMP4+ prior to dissociation. Day 45 cells were stained for Myosin heavy chain (MYH1), which is expressed in myocytes, myotubes and skeletal myofibers. (B) Day 45 cells derived from Somitoids were fixed and stained for Myosin heavy chain (MYH1) and Hoechst (nuclear stain). Somite-stage organoids (Day 5) were treated with WNT+/SHH-/BMP4+ signaling modulators for 48h from day 5 to day 7. (C) Same as in (B) but with control cells that were not pulse-treated with WNT+/SHH-/BMP4+ signaling modulators but instead cultured in basal media from day 5 to day 7.

## Supplemental Files

**Supplemental file 1:** Supplemental Table 1, cluster-based marker genes of scRNA-seq dataset

**Supplemental file 2:** Supplemental Table 2, RT-qPCR primer sequences

**Figure 3-Video 1**, confocal z-stacks of Somitoids and control organoids immunostained for PAX3 and NCAD showing *in vitro* somites

## Source Data Files

Fig 1-Source Data 1: qPCR raw data of PM organoid differentiation.

Fig 2-Source Data 1: Quantification of organoid diameter from primary screen.

Fig 2-Source Data 2: Quantification of average PAX3 levels per organoid from primary screen.

Fig 3-Source Data 1: Quantification of average PAX3 levels per organoid from secondary screen.

Fig 3-Source Data 2: Somite quantification of secondary screen.

Fig 5-Source Data 1: Comparative somite size quantification of *in vitro* somites and human somites from the Carnegie collection.

Fig 5-Source Data 2: qPCR raw data of sclerotome differentiation.

Fig 3-Supplemental Fig 2-Source Data 1: qPCR data of calculated expression levels of selected treatment regimes from secondary screen.

Fig 3-Supplemental Fig 3-Source Data 1: Somite quantification of technical replicates using the NCRM1 cell line.

Fig 3-Supplemental Fig 3-Source Data 2: Somite quantification of biological replicates using the NCRM1, ACTB-GFP, and WTC cell lines.

## References

1. Aulehla A, Wehrle C, Brand-Saberi B, Kemler R, Gossler A, Kanzler B, Herrmann BG. 2003. Wnt3a plays a major role in the segmentation clock controlling somitogenesis. Dev Cell 4:395–406. doi:10.1016/S1534-5807(03)00055-8

2. Aulehla A, Wiegraebe W, Baubet V, Wahl MB, Deng C, Taketo M, Lewandoski M, Pourquié O. 2008. A beta-catenin gradient links the clock and wavefront systems in mouse embryo segmentation. Nat Cell Biol 10:186–193. doi:10.1038/ncb1679

3. Beccari L, Moris N, Girgin M, Turner DA, Baillie-Johnson P, Cossy A-C, Lutolf MP, Duboule D, Arias AM. 2018. Multi-axial self-organization properties of mouse embryonic stem cells into gastruloids. Nature 562:272–276.

4. Chal J, Oginuma M, Al Tanoury Z, Gobert B, Sumara O, Hick A, Bousson F, Zidouni Y, Mursch C, Moncuquet P, Tassy O, Vincent S, Miyanari A, Bera A, Garnier J-M, Guevara G, Hestin M, Kennedy L, Hayashi S, Drayton B, Cherrier T, Gayraud-Morel B, Gussoni E, Relaix F, Tajbakhsh S, Pourquié O. 2015. Differentiation of pluripotent stem cells to muscle fiber to model Duchenne muscular dystrophy. Nat Biotechnol 33:962–969. doi:10.1038/nbt.3297

5. Chal J, Pourquié O. 2017. Making muscle: skeletal myogenesis *in vivo* and *in vitro*. Development 144:2104–2122.

6. Chu L-F, Mamott D, Ni Z, Bacher R, Liu C, Swanson S, Kendziorski C, Stewart R, Thomson JA. 2019. An In Vitro Human Segmentation Clock Model Derived from Embryonic Stem Cells. Cell Rep 28:2247–2255.e5. doi:10.1016/j.celrep.2019.07.090

7. Delfini M-C, Dubrulle J, Malapert P, Chal J, Pourquié O. 2005. Control of the segmentation process by graded MAPK/ERK activation in the chick embryo. Proceedings of the National Academy of Sciences 102:11343–11348. doi:10.1073/pnas.0502933102

8. Dequeant ML, Glynn E, Gaudenz K, Wahl M, Chen J, Mushegian A, Pourquie O. 2006. A Complex Oscillating Network of Signaling Genes Underlies the Mouse Segmentation Clock. Science 314:1595–1598. doi:10.1126/science.1133141

9. Diaz-Cuadros M, Wagner DE, Budjan C, Hubaud A, Tarazona OA, Donelly S, Michaut A, Al Tanoury Z, Yoshioka-Kobayashi K, Niino Y, Kageyama R, Miyawaki A, Touboul J, Pourquié O. 2020. In vitro characterization of the human segmentation clock. Nature 580:113–118. doi:10.1038/s41586-019-1885-9

10. Dubrulle J, McGrew MJ, Pourquié O. 2001. FGF Signaling Controls Somite Boundary Position and Regulates Segmentation Clock Control of Spatiotemporal Hox Gene Activation. Cell 106:219–232. doi:10.1016/S0092-8674(01)00437-8

11. Dunty WC Jr, Biris KK, Chalamalasetty RB, Taketo MM, Lewandoski M, Yamaguchi TP. 2008. Wnt3a/beta-catenin signaling controls posterior body development by coordinating mesoderm formation and segmentation. Development 135:85–94. doi:10.1242/dev.009266

12. Fan CM, Porter JA, Chiang C, Chang DT, Beachy PA, Tessier-Lavigne M. 1995. Long-range sclerotome induction by sonic hedgehog: direct role of the amino-terminal cleavage product and modulation by the cyclic AMP signaling pathway. Cell 81:457– 465. doi:10.1016/0092-8674(95)90398-4

13. Fan CM, Tessier-Lavigne M. 1994. Patterning of mammalian somites by surface ectoderm and notochord: evidence for sclerotome induction by a hedgehog homolog. Cell 79:1175–1186. doi:10.1016/0092-8674(94)90009-4

14. Freedman BS, Brooks CR, Lam AQ, Fu H, Morizane R, Agrawal V, Saad AF, Li MK, Hughes MR, Werff RV, Peters DT, Lu J, Baccei A, Siedlecki AM, Valerius MT, Musunuru K, McNagny KM, Steinman TI, Zhou J, Lerou PH, Bonventre JV. 2015. Modelling kidney disease with CRISPR-mutant kidney organoids derived from human pluripotent epiblast spheroids. Nat Commun 6:8715. doi:10.1038/ncomms9715

15. Greco TL, Takada S, Newhouse MM, McMahon JA, McMahon AP, Camper SA. 1996. Analysis of the vestigial tail mutation demonstrates that Wnt-3a gene dosage regulates mouse axial development. Genes Dev 10:313–324. doi:10.1101/gad.10.3.313

16. Hafemeister C, Satija R. 2019. Normalization and variance stabilization of single-cell RNA-seq data using regularized negative binomial regression. bioRxiv. doi:10.1101/576827

17. Hubaud A, Pourquié O. 2014. Signalling dynamics in vertebrate segmentation. Nat Rev Mol Cell Biol 15:709–721.

18. Li CH, Tam PKS. 1998. An iterative algorithm for minimum cross entropy thresholding. Pattern Recognit Lett 19:771–776. doi:10.1016/S0167-8655(98)00057-9

19. Loh KM, Chen A, Koh PW, Deng TZ, Sinha R, Tsai JM, Barkal AA, Shen KY, Jain R, Morganti RM, Shyh-Chang N, Fernhoff NB, George BM, Wernig G, Salomon REA, Chen Z, Vogel H, Epstein JA, Kundaje A, Talbot WS, Beachy PA, Ang LT, Weissman IL. 2016. Mapping the Pairwise Choices Leading from Pluripotency to Human Bone, Heart, and Other Mesoderm Cell Types. Cell 166:451–467. doi:10.1016/j.cell.2016.06.011

20. Manfrin A, Tabata Y, Paquet ER, Vuaridel AR, Rivest FR, Naef F, Lutolf MP. 2019. Engineered signaling centers for the spatially controlled patterning of human pluripotent stem cells. Nat Methods 16:640–648. doi:10.1038/s41592-019-0455-2

21. Matsuda M, Yamanaka Y, Uemura M, Osawa M, Saito MK, Nagahashi A, Nishio M, Guo L, Ikegawa S, Sakurai S, Kihara S, Maurissen TL, Nakamura M, Matsumoto T, Yoshitomi H, Ikeya M, Kawakami N, Yamamoto T, Woltjen K, Ebisuya M, Toguchida J, Alev C. 2020. Recapitulating the human segmentation clock with pluripotent stem cells. Nature 580:124–129. doi:10.1038/s41586-020-2144-9

22. Moris N, Anlas K, van den Brink SC, Alemany A, Schröder J, Ghimire S, Balayo T, van Oudenaarden A, Martinez Arias A. 2020. An in vitro model of early anteroposterior organization during human development. Nature 582:410–415. doi:10.1038/s41586-020-2383-9

23. Oates AC, Morelli LG, Ares S. 2012. Patterning embryos with oscillations: structure, function and dynamics of the vertebrate segmentation clock. Development 139:625– 639. doi:10.1242/dev.063735

24. Palla A, Blau H. 2020. The clock that controls spine development modelled in a dish. Nature 580:32–34. doi:10.1038/d41586-020-00322-y

25. Sakurai H, Sakaguchi Y, Shoji E, Nishino T, Maki I, Sakai H, Hanaoka K, Kakizuka A, Sehara-Fujisawa A. 2012. In vitro modeling of paraxial mesodermal progenitors derived from induced pluripotent stem cells. PLoS One 7:e47078. doi:10.1371/journal.pone.0047078

26. Schindelin J, Arganda-Carreras I, Frise E, Kaynig V, Longair M, Pietzsch T, Preibisch S, Rueden C, Saalfeld S, Schmid B, Tinevez J-Y, White DJ, Hartenstein V, Eliceiri K, Tomancak P, Cardona A. 2012. Fiji: an open-source platform for biological-image analysis. Nat Methods 9:676–682. doi:10.1038/nmeth.2019

27. Stuart T, Butler A, Hoffman P, Hafemeister C, Papalexi E, Mauck WM, Hao Y, Stoeckius M, Smibert P, Satija R. 2019. Comprehensive Integration of Single-Cell Data. Cell 0.

28. Tan JY, Sriram G, Rufaihah AJ, Neoh KG, Cao T. 2013. Efficient derivation of lateral plate and paraxial mesoderm subtypes from human embryonic stem cells through GSKi-mediated differentiation. Stem Cells Dev 22:1893–1906.

29. Tonegawa A, Funayama N, Ueno N, Takahashi Y. 1997. Mesodermal subdivision along the mediolateral axis in chicken controlled by different concentrations of BMP-4. Development 124:1975–1984.

30. Traag VA, Waltman L, van Eck NJ. 2019. From Louvain to Leiden: guaranteeing well-connected communities. Sci Rep 9:5233. doi:10.1038/s41598-019-41695-z

31. Tzouanacou E, Wegener A, Wymeersch FJ, Wilson V, Nicolas J-F. 2009. Redefining the progression of lineage segregations during mammalian embryogenesis by clonal analysis. Dev Cell 17:365–376.

32. van den Brink SC, Baillie-Johnson P, Balayo T, Hadjantonakis A-K, Nowotschin S, Turner DA, Martinez Arias A. 2014. Symmetry breaking, germ layer specification and axial organisation in aggregates of mouse embryonic stem cells. Development 141:4231–4242. doi:10.1242/dev.113001

33. Veenvliet JV, Bolondi A, Kretzmer H, Haut L, Scholze-Wittler M, Schifferl D, Koch F, Pustet M, Heimann S, Buschow R, Wittler L, Timmermann B, Meissner A, Herrmann BG. 2020. Mouse embryonic stem cells self-organize into trunk-like structures with neural tube and somites. bioRxiv. doi:10.1101/2020.03.04.974949

34. Xi H, Fujiwara W, Gonzalez K, Jan M, Liebscher S, Van Handel B, Schenke-Layland K, Pyle AD. 2017. In Vivo Human Somitogenesis Guides Somite Development from hPSCs. CellReports 18:1573–1585.

35. Yamaguchi TP, Takada S, Yoshikawa Y, Wu N, McMahon AP. 1999. T (Brachyury) is a direct target of Wnt3a during paraxial mesoderm specification. Genes & Development. doi:10.1101/gad.13.24.3185

